# CellChem: Cellular transcriptional responses reshape molecular representation space for efficient and multi-scale drug discovery

**DOI:** 10.64898/2026.04.22.719826

**Authors:** Jiaxiao Chen, Letian Lin, Yishen Wang, Yifan Lin, Yu Li, Weilin Zhang, Yiwei Fu, Jin Xie, Jintao Zhu, Chao Sun, Guqin Shi, Su Qian, Zheng Wang, Haoyu Lin, Liying Wang, Minghua Deng, Luhua Lai, Jianfeng Pei

## Abstract

Despite decades of progress in computational drug discovery, deep learning-based molecular representation models remain largely structure-centric, assuming that chemical similarity approximates functional similarity. However, drug effects in cells are shaped not only by chemical similarity but also by molecular interactions in the cellular context. To capture this complexity, we introduce CellChem, a cellular-chemical representation-learning framework that learns cell-guided molecular representations of small-molecule actions within cells. By incorporating large-scale cellular transcriptional profiles during pretraining, CellChem reshapes molecular representation space from structure-centric to a balanced integration of structure and function. The learned CellChem molecular representations exhibit biologically meaningful geometric organization, such that distances between molecules encode not only structural similarity but also similarity in the cellular responses they elicit, independent of downstream tasks. Using downstream derivative models such as cell-guided compound-protein interaction prediction and drug-induced transcriptional response profile generation, CellChem supports highly efficient, multi-scale drug discovery, achieving significantly better performance than traditional models.

## Introduction

Molecular representation learning has recently improved by focusing on molecular structure. Using large-scale training data, these methods learn embeddings that capture the chemical patterns encoded in molecular graphs, such as local functional groups, global scaffolds, and more complex structural features, providing strong, transferable, structure-aware features for a wide range of downstream tasks. For example, MolCLR leverages vast unlabeled datasets and a graph neural network encoder to construct molecular embeddings that accurately describe structural similarity and chemical organization^1^. Other approaches improve structural fidelity by pretraining with three-dimensional conformations^3^ or by integrating multiple structure-based inputs to capture complementary geometric and physicochemical cues^2-6^. These strategies assume that chemical similarity approximates functional similarity. However, we can now use massive cellular transcriptome data sets^7^ to improve molecular representations.

Cells respond to many compounds by changing their transcriptional programs and regulatory networks, and these changes are influenced by factors such as dose, treatment duration, and pathway dynamics^8-10^. Structure-based representations alone do not capture this complexity, because 1) structurally dissimilar compounds may have similar functions and 2) structurally similar compounds may have dissimilar functions: for example, the well-documented phenomenon of activity cliffs^11, 12^. Thus, structure-based pretraining alone cannot elucidate mechanisms of action or drug behavior in cells. Therefore, research using virtual cells is driving a shift from predicting molecular structure and properties to understanding molecular behaviors and cellular effects^13, 14^. Large single-cell foundation models provide representations that can encode phenotype information^15-17^.

Here, we present CellChem, a training system that uses cellular transcriptional response data to shape the geometry of molecular representation space. Previous systems have used cell-state information to guide downstream drug-discovery tasks, including predicting or generating perturbation profiles from chemical structures or designing molecules to match transcriptomic signatures ^18-23^. However, these systems do not integrate cell responses into the core molecular representation; they serve only as a downstream conditioning signal. Here, we combine chemical space with cell response space, using the cell transcriptional response to construct a new molecular representation space that incorporates both chemical-structure and cellular-functional similarity.

CellChem not only encodes structural similarity but also creates embeddings that organize molecules based on functional relationships, independent of downstream tasks. This produces a representation that generalizes across different tasks and reflects underlying cellular mechanisms. Thus, molecular similarity depends not only on chemical structure alone but also on shared effects in cellular space and functional organization. Our new molecular representation paradigm should be universal, reusable across tasks, and interpretable for multi-scale drug discovery with cell-guided enhancement. By integrating cross-modal signals into an enhanced, unified knowledge framework, we expect CellChem to outperform previous downstream conditioning methods.

We used CellChem to integrate functional information into downstream tasks, thereby coherently integrating molecular structure, cellular response, and biological function. We demonstrated that CellChem enhanced scaffold-level generalization in compound-protein interaction (CPI) prediction and enabled accurate generation of cellular transcriptional profiles for unseen chemical scaffolds. In a drug screening campaign, combining CPI-based ranking with gene expression profile matching accelerated PLK1 inhibitor discovery and improved screening performance. We next addressed a more challenging application involving the orphan GPCR G Protein-Coupled Receptor 6 (GPR6), for which no transcriptional profile is available. By combining CPI-based ranking with model-predicted cellular transcriptional profiles, we identified inverse agonists with novel scaffolds. CellChem-generated cellular transcriptional profiles not only provide cellular-level representations of molecular perturbations for virtual screening but also offer gene-level functional readouts. We demonstrated the system-level effects of traditional Chinese medicinal compounds whose predicted transcriptomic responses accurately explained their reported mechanisms, pathway activities, and gene-regulatory programs. Thus, CellChem supports highly efficient, multi-scale drug discovery.

## Results

### Overview of CellChem

As a cross-modal cellular-chemical framework, CellChem integrates data on molecular structure and cellular transcriptional response. The molecular dataset comprises 10 million compounds from the PubChem database^1^, the cellular transcriptional response data come from the LINCS database^7^, and gene expression profiles are derived from the Connectivity Map (CMAP)—the large-scale molecular perturbation expression repository. This database uses L1000 high-throughput gene expression technology to generate comprehensive expression profiles via sequencing methods analogous to those used in conventional approaches. We used LINCS Level 5 signatures, which represent consensus gene expression profiles computed from multiple biological replicates responding to the same perturbation after normalization and replicate-level quality control. The final curated perturbation dataset comprised 505,052 entries (Supplementary Table S1); statistical analyses are provided in Supplementary Figure S1.

CellChem uses a four-stage pretraining strategy that integrates functional information into downstream tasks and establishes a unified framework that coherently links molecular structure, cellular response, and biological function (Figure 1a). During stage 1 (Figure 1b), contrastive learning was applied with random masking of 25% of atoms to create augmented molecular views, following the settings of the established molecule contrastive pretraining practice^24^. The molecular encoder was initially pretrained independently using a Graph Transformer architecture^25^ to learn structure-invariant chemical features. This pretraining embedding produced chemically coherent latent representations that serve as the structural foundation for subsequent alignment.

**Figure 1:**
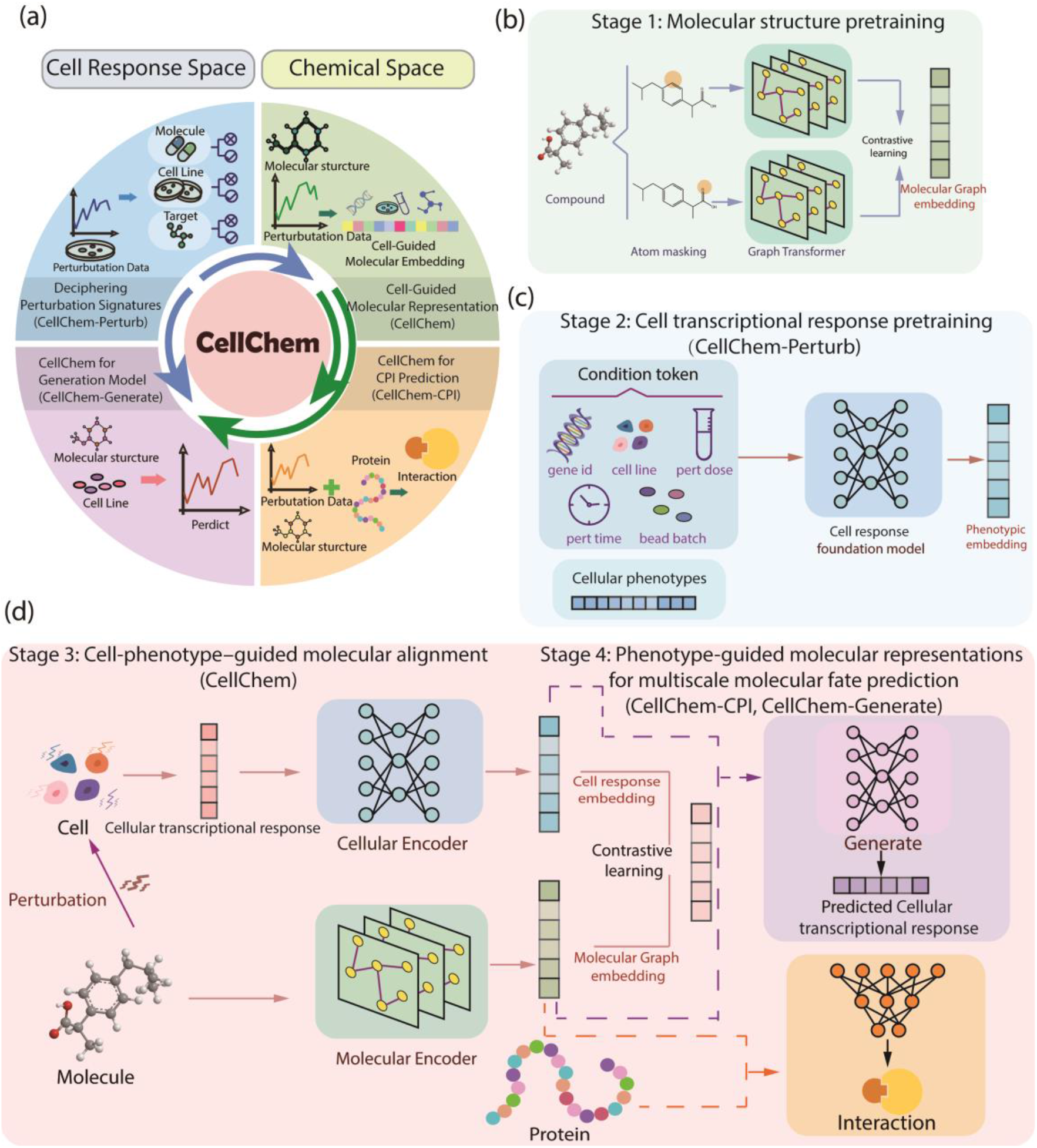
Overview of the CellChem framework for cellular transcriptional response–guided molecular representation learning. (a) Conceptual schematic of CellChem bridging the chemical space and cellular transcriptional response space. Molecular structures and their paired cellular transcriptional response profiles are the inputs for learning cell-guided molecular representations. These extend structure-based similarity to a functional and cell-state organization. (b) Stage 1: Molecular structure pretraining. A graph transformer is pretrained on large-scale unlabeled compounds using atom masking and contrastive learning to produce molecular graph embeddings. (c) Stage 2: Cellular transcriptional response pretraining. A cell response foundation model is pretrained to encode transcriptional responses from perturbation data, conditioned on metadata tokens, yielding cell response embeddings.(d) Stage 3–4: Cell-guided molecular alignment and downstream usage. Stage 3 aligns molecular graph embeddings with cellular transcriptional response embeddings via contrastive learning, reshaping the molecular embedding geometry into a functionally organized manifold. Stage 4: multi-scale drug discovery tasks, such as predicting cellular transcriptional response to a molecule and compound–protein interaction (CPI), use the shared latent space to link macroscale cellular response modeling with microscale molecular interaction mechanisms.

In stage 2 (Figure 1c), we pretrained a cellular encoder (CellChem-Perturb) on large-scale perturbational transcriptomic profiles by adapting an RNA-seq foundation model^24^ to the perturbation setting. This encoder was further conditioned on experimental variables that modulate cellular transcriptional responses, implemented using condition tokens that encode variables such as cell line, dose, and treatment duration (Table S1 provides detailed information on cellular transcriptional responses). Through cellular perturbation pretraining, the encoder organizes transcriptional response embeddings into a biologically relevant latent space, independent of structural information.

In stage 3 (Figure 1d, left), CellChem performs cross-modal alignment, using cellular data to transform the latent molecular embedding space. Rather than serving as prediction targets, transcriptional responses guide the organization of molecular representations, embedding functional cellular information into the latent space. Latent representations of each molecule and its corresponding cellular transcriptional response are treated as positive pairs, whereas mismatched combinations in the same batch serve as negative pairs. A contrastive loss function pulls structurally dissimilar but phenotypically consistent molecules closer and pushes those with phenotypically divergent molecules further apart.

In stage 4 (Figure 1d, right), cell-guided molecular representations supported both compound– protein interaction prediction (CellChem-CPI) and the generation of cellular transcriptional profiles (CellChem-Generate). The four-stage pretraining enabled CellChem to outperform task-specific models in Stage 4, moving toward a general-purpose, function-aware representation of molecules guided by cellular context. Detailed architectural specifications and implementation settings for all the training are provided in the Methods.

### Cellular transcriptional responses redefine molecular representation geometry

CellChem uses contrastive learning on cellular transcriptional responses to reshape the geometry of molecular representations into a cell-aware space. Deep learning-based modeling of these data captures more biological information than conventional single-cell analysis pipelines^26, 27^. Transcriptional response embeddings provide the functional basis for transforming molecular representations from a structure-only space to a cell-guided space.

The cellular encoder (CellChem-Perturb) first learns the biologically meaningful organization across molecular, target, and cell-line levels, removing batch effects and establishing a transcriptomic latent space that encodes functional and mechanistic information. CellChem-Perturb initially produces cellular transcriptional response embeddings with batch-dependent separation during early training stages (epoch 5), which progressively converge into a well-integrated manifold by epoch 88, indicating suppression of technical variation and preservation of pharmacologically relevant signals (Supplementary Figure 2a–c). Therefore, CellChem-Perturb outperformed two widely used association-based frameworks, L1000FWD^26^ and SigCom LINCS^27^, in extracting structure from noisy perturbation data across molecule-, target-, and cell–line-level identification tasks (Figure 2a). Implementation details and dimensionality reduction procedures for all three methods are provided in the Methods section. CellChem-Perturb achieved higher Adjusted Rand Index (ARI) and Fowlkes–Mallows Index (FMI) scores than L1000FWD and SigCom LINCS for multiple compounds, including mirdametinib (LINCS name: PD-0325901) ^28^ and tanespimycin ^29^ (Figure 2b), demonstrating the value of deep-learning-based perturbation profiles encoding in capturing biologically relevant cellular structure. UMAP^30^ projections illustrate this multi-scale organization: at the target level, CellChem-Perturb demonstrated increased inter-cluster separation (Figure 2a, middle), whereas at the cell-line level, it produced sharper boundaries between cellular contexts (Figure 2a, right) with improved clustering metrics. We summarize the gains across molecular, target, and cell-line dimensions in Figure 2b.

**Figure 2:**
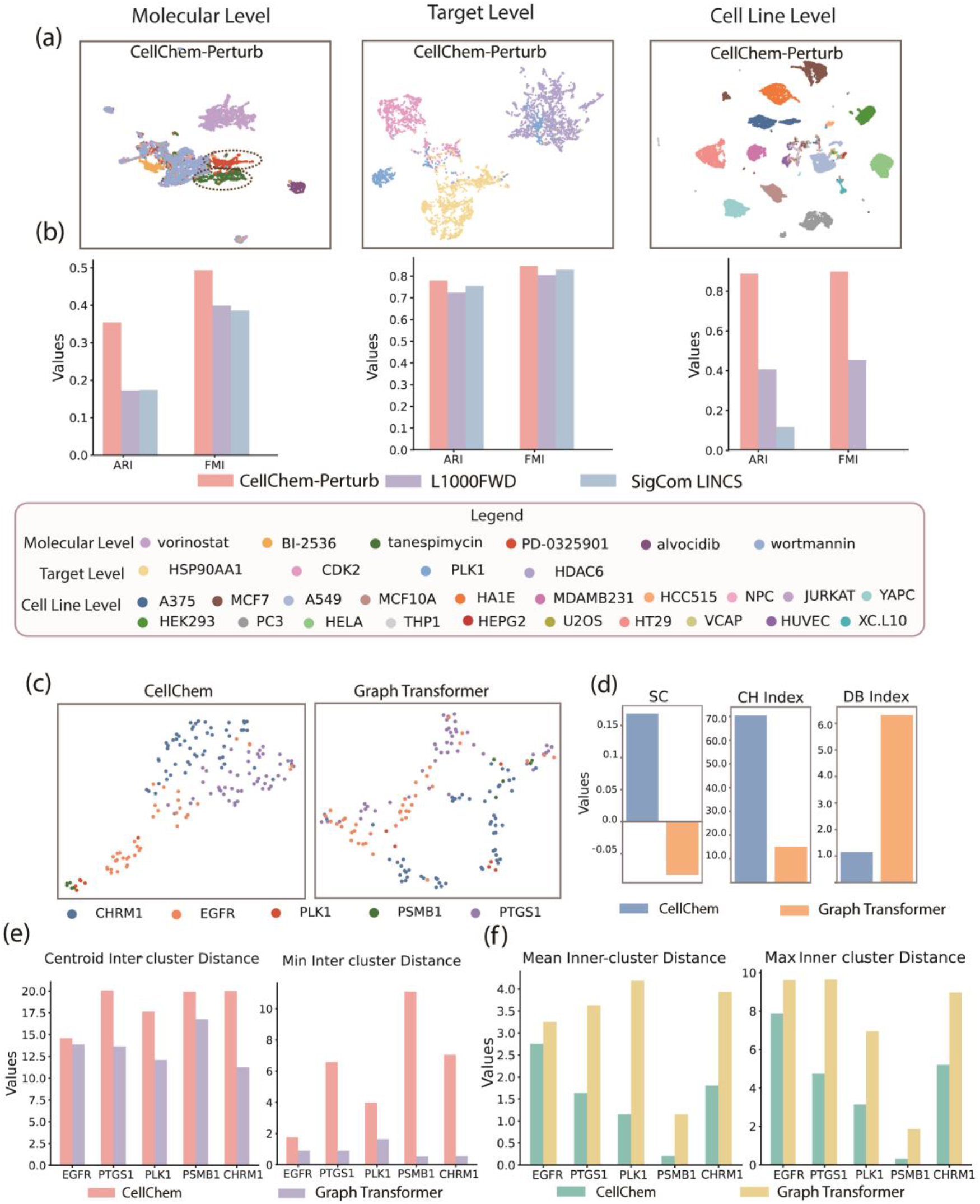
Cell-guided molecular embeddings (CellChem) capture functional organization. (a) Visualization of perturbation features via UMAP projection: molecular-level, target-level, and cell-line-level analyses. (b) Quantitative comparison of CellChem vs. L1000FWD and SigCom LINCS using Adjusted Rand Index (ARI) and Fowlkes–Mallows Index (FMI) for all three evaluation tiers. (c) Latent space visualization comparison. Left: UMAP projection of the top 5 target molecules using the cell-guided CellChem model. Right: UMAP projection of the same molecules using Graph Transformer (molecular structure data only). (d) Quantitative evaluation of clustering performance. Comparison between CellChem and Graph Transformer without cellular transcriptional responses guidance using three clustering metrics: Silhouette Coefficient (SC), Calinski-Harabasz Index (CH index), and Davies–Bouldin Index (DB index). (e) Inter-class distance analysis. Statistical comparison of inter-class distances (Euclidean distance) in the latent space between the two models. (f) Intra-class compactness analysis. Statistical comparison of intra-class distances, measured as the mean Euclidean distance from samples to their respective cluster centroids.

This high-quality cellular embedding provided the foundation for cell-guided molecular representation learning. Unlike a structure-only Graph Transformer–pretrained molecular embedding model, which organizes compounds primarily by chemical scaffold similarity, CellChem incorporates cellular transcriptomic response as a functional prior. Compared with the baseline molecular structure–pretrained Graph Transformer, the cell-guided molecular encoder (CellChem) learned by CellChem-Perturb induced target-specific functional neighborhoods in molecular space. Proteasome subunit beta 1 (PSMB1)-targeting compounds^31^, which are widely dispersed in structure-only embeddings due to scaffold diversity (Figure 2c, right), collapsed into compact clusters after cellular alignment (Figure 2c, left). Similarly, PTK1S-targeting molecules^32^ transitioned from fragmented, scaffold-driven subgroups in the Graph Transformer baseline into a single manifold organized by a shared mechanism (Figure 2c, left). A quantitative evaluation of many compounds confirmed this geometric reorganization (Figure 2d). The silhouette coefficient (SC) increased from −0.082 to 0.1683, the Calinski-Harabasz index increased from 15.13 to 70.39, and the Davies–Bouldin index decreased from 6.308 to 1.155. A geometric analysis showed that inter-target distances doubled, whereas intra-target molecular dispersion decreases by 55% (Figure 2e–f), indicating expansion of functional boundaries and compaction of mechanistic cores. Notably, these improvements surpassed structure-only Graph Transformer pretraining, which does not use cellular response supervision to align molecules by shared functional mechanisms across scaffolds. Thus, the transcriptional response encoded by CellChem-Perturb provided the biological and mechanistic foundation for CellChem to reshape molecular representations into functionally organized embedding spaces, grouping molecules by their cellular effects rather than by chemical similarity alone.

### CellChem for Compound-Protein Interaction Prediction

CPI prediction determines whether a compound and a target protein interact. Deep learning CPI models take protein sequences and integrate them with compound structures to train end-to-end interaction predictors. For example, GraphBAN^33^ learns compound–protein representations from molecular graphs using graph neural networks and knowledge distillation, and TransformerCPI^34^ models both compounds and proteins as token sequences, while its embeddings remain largely structure-driven. CPI can screen vast, largely uncharacterized libraries of chemicals with novel scaffolds that are distant from known active ligands. Here, cellular transcriptional profiles provided a complementary, mechanism-linked signal that is not tied to scaffold similarities. We integrated transcriptional profiles as a functional prior to enhance CPI prediction (Figure 3a). Detailed network specifications for the model are provided in Supplementary Figure 4a (an expanded view of Figure 3a) and in the Methods. CellChem-CPI is built in chemical space and provides inference without transcriptional profiles, using the cell-guided molecular embedding learned by CellChem.

**Figure 3:**
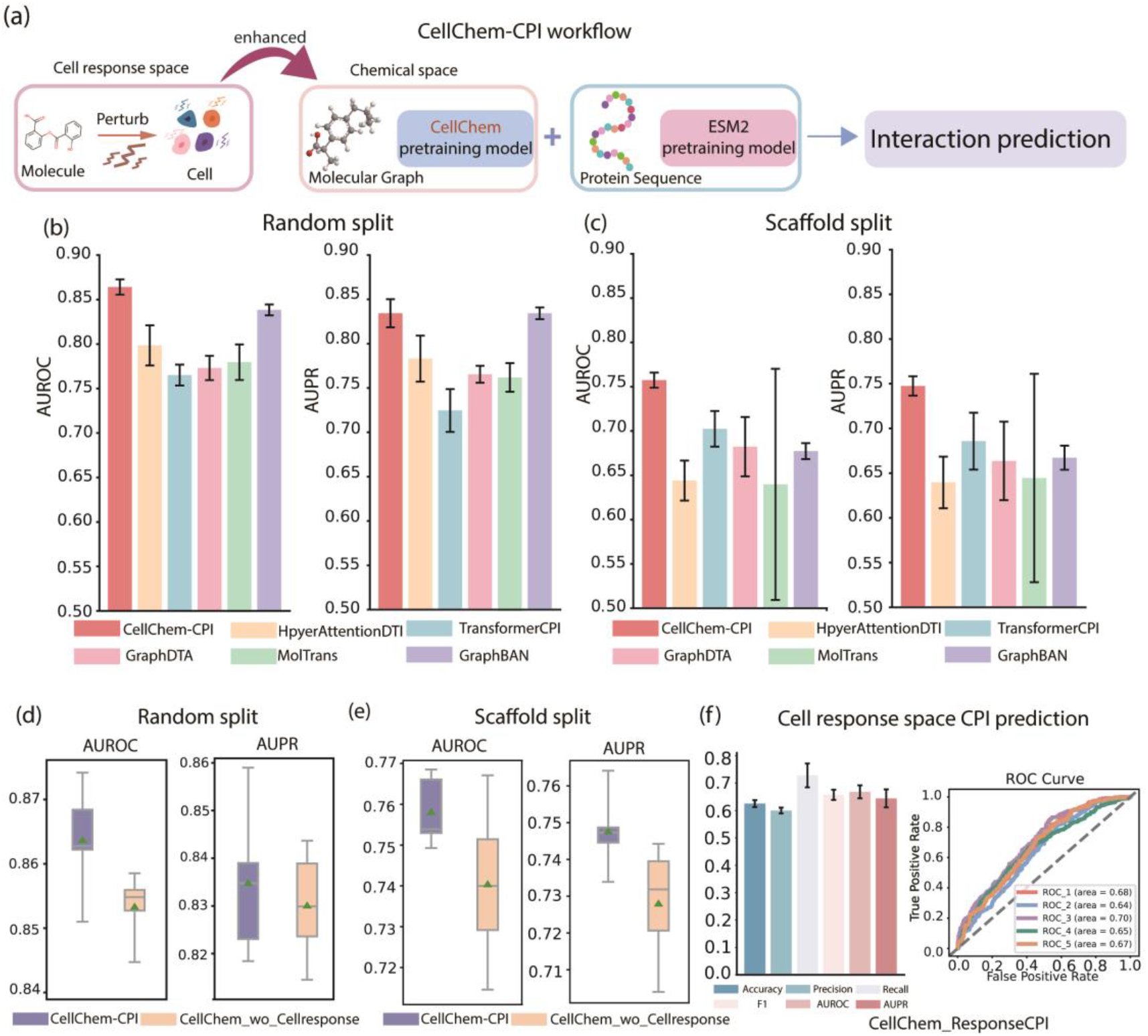
Performance evaluation of CellChem-CPI prediction. (a) Architecture of CPI prediction model integrating protein sequence features, molecular structures, and cellular transcriptional response data. The complete computational workflow from raw data input to final prediction output is shown in greater detail in Supplementary Figure 4a. (b) (c) Systematic comparison results between the CellChem model and sequence-based prediction methods on randomly-split and scaffold-split datasets. Error bars represent the standard deviation from five-fold cross-validation. (d) (e) Model architecture ablation study. Controlled comparison of 1) cell-pretrained CellChem for CellChem-CPI and 2) CellChem_wo_Cellresponse using molecular structural features only. (f) Prediction performance using cellular transcriptional response alone. The model demonstrated significant discriminative power as the area under the curve (AUC = 0.6683 ± 0.026), confirming that cellular transcriptional response is an effective auxiliary feature for CPI prediction. Receiver operating characteristic (ROC) curve analysis using transcriptional response alone. The AUC reached 0.70 on the test set, indicating good predictive performance.

We compared CellChem-CPI against structure-only CPI methods^33-37^ with both random-split and scaffold-split protocols. The dataset was constructed from the intersection of CMAP^7^ and DrugBank^38^. In the random-split setting, CellChem-CPI achieved an area under the receiver operating characteristic curve (AUROC) of 0.864, outperforming the best structure-only CPI methods (Figure 3b). In the scaffold-split setting, CellChem-CPI achieved even better results, with AUROC and area under the precision-recall curve (AUPRC) of 0.757 and 0.747, respectively, compared with 0.702 and 0.686 for the best structure-only method (Figure 3c). In an ablation study, we showed that this improvement did not result from the new model architecture; when we removed the cell-guided molecular embeddings while keeping the same CPI model architecture (CellChem_wo_Cellresponse), the model’s performance decreased, especially for the scaffold split (Figure 3d, e). Therefore, the improved performance results from the cell-guided representation geometry in CellChem, which provides additional functional information, thereby strengthening CPI inference when extrapolating to novel chemical scaffolds. We also performed the reciprocal experiment by removing structural information and using only transcriptional response signals. We used CRISPR gene knockouts to provide a functional representation of proteins by capturing the cell-wide transcriptional effects of removing a protein. We combine these embeddings with compound-induced transcriptional profiles to study functional relationships in a cell response–based space, independent of protein structure. In this cell response space, the average Pearson correlation was 0.1552 for molecular profiles and 0.2062 for protein knockout profiles (Supplementary Figure 5), indicating that molecule-induced and protein-induced perturbations occupied a coherently organized space rather than random noise. Using only these cellular response profiles, CellChem-ResponseCPI (Supplementary Figure 4b) accurately matched compounds to their interacting proteins, achieving AUROC = 0.67 and AUPR = 0.65 under 5-fold validation (Figure 3f). However, CellChem-CPI, which integrates both cell response space and chemical space, outperformed CellChem-ResponseCPI, which operates only in cell response space, and CellChem_wo_Cellresponse, which relies only on chemical space.

### PLK1 inhibitor discovery by CellChem molecular embeddings and cell response

We screened for drugs against PLK1, a canonical cell cycle kinase targeted by inhibitors, using CellChem-CPI and CellChem-Perturb^39-41^. CellChem-CPI prioritizes interactions, ranking compounds for predicted PLK1 engagement, thereby narrowing a large library to a manageable, focused set. In parallel, CellChem-Perturb prioritizes cellular responses by embedding the cellular transcriptional response to compounds, ranking compounds according to their Pearson similarity to the positive control compound NMS-1286937. We also evaluated the screening performance of CellChem-CPI, CellChem-Perturb, and Glide SP docking^42, 43^, comparing them to the integrated strategy. We tested the performance of CellChem on a reference set of 29 PLK1-active compounds curated from ChEMBL in the LINCS collection^44^ using CellChem-Perturb + CellChem-CPI, CellChem-CPI, CellChem-Perturb, and Glide docking (Figure 4a). CellChem-CPI showed strong rankings for known PLK1 inhibitors and substantially outperformed Glide docking; therefore, functional compatibility was more predictive than structural fit. Similarly, the CellChem-Perturb ranking favored compounds whose transcriptional response embeddings were consistent with PLK1 inhibition. The integrated CellChem-Perturb + CellChem-CPI strategy best identified known active compounds, achieving an AUC of 0.91, compared to 0.85 for CellChem-Perturb, 0.81 for CellChem-CPI, and 0.57 for Glide docking rankings (Figure 4b). The enrichment factor curves showed that integrated CellChem-based methods were stronger in the early enrichment stage (top 5%) than Glide, and the integrated strategy yielded the most robust prioritization of known active compounds in the top-ranked fraction (Figure 4c). Based on kernel density distributions (Figure 4d), CellChem clearly separated active and inactive compounds, and the bimodal distribution among nominally inactive molecules suggests that this group may contain undiscovered PLK1 inhibitors. Compared to structure-based CPI methods, CellChem-CPI achieved superior enrichment in PLK1 inhibitor screening (Figure 4e). Overall, the integrated CellChem-Perturb + CellChem-CPI strategy achieved the best enrichment performance for screening PLK1 inhibitors.

**Figure 4:**
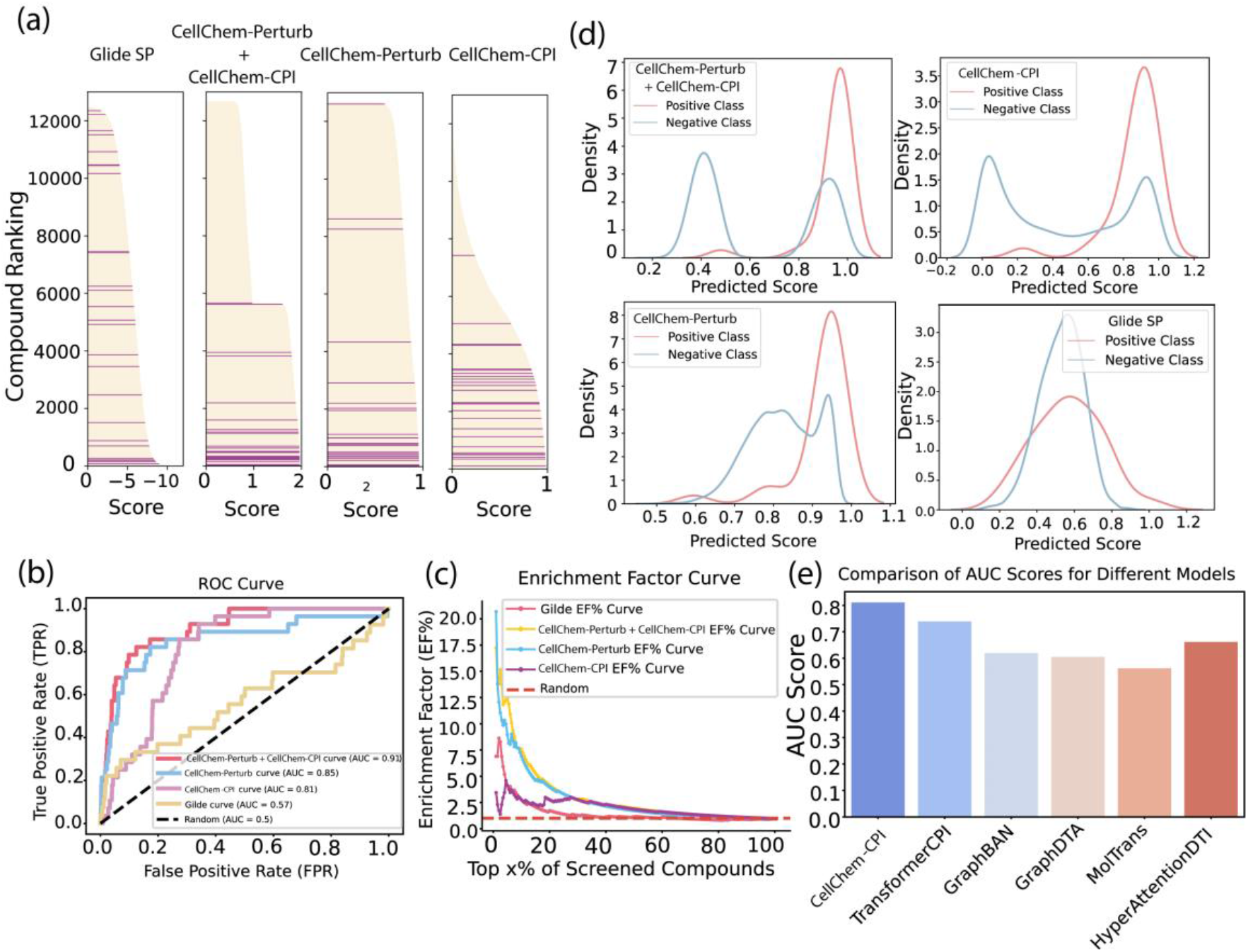
Performance evaluation of CellChem for drug screening using CellChem-perturb and CellChem-CPI model. (a) Ranking plot of PLK1 inhibitors by four strategies (CellChem-perturb+CellChem-CPI, CellChem-CPI, CellChem-perturb, and Glide docking). The ranking positions of known active compounds are shown as purple lines. (b) Receiver operating characteristic (ROC) curve comparison for four screening methods (CellChem-perturb+CellChem-CPI, CellChem-CPI, CellChem-perturb, and Glide docking) on the test set. CellChem-perturb+CellChem-CPI demonstrated the best overall discriminative performance, with an area under the curve (AUC)=0.91. (c) Comparison of enrichment factor curve for the four methods across 1%–100% hit molecules. (d) Kernel density distribution curves of active and inactive molecules for the four methods. (e) Quantitative AUC comparisons of four sequence-based deep learning methods.

We then tested high-scoring compounds experimentally (Figure 4b upper left) to determine whether CellChem could identify previously unannotated PLK1 inhibitors. The workflow (Figure 5a) began by excluding known active molecules and yielded 70 top-ranked commercially available compounds for laboratory validation. We determined that 28 compounds at 50 μM showed >50% PLK1 inhibition—a hit rate of approximately 40% (Figure 5b). This success rate is higher than is typical for structure-based virtual screening alone^45^.

**Figure 5:**
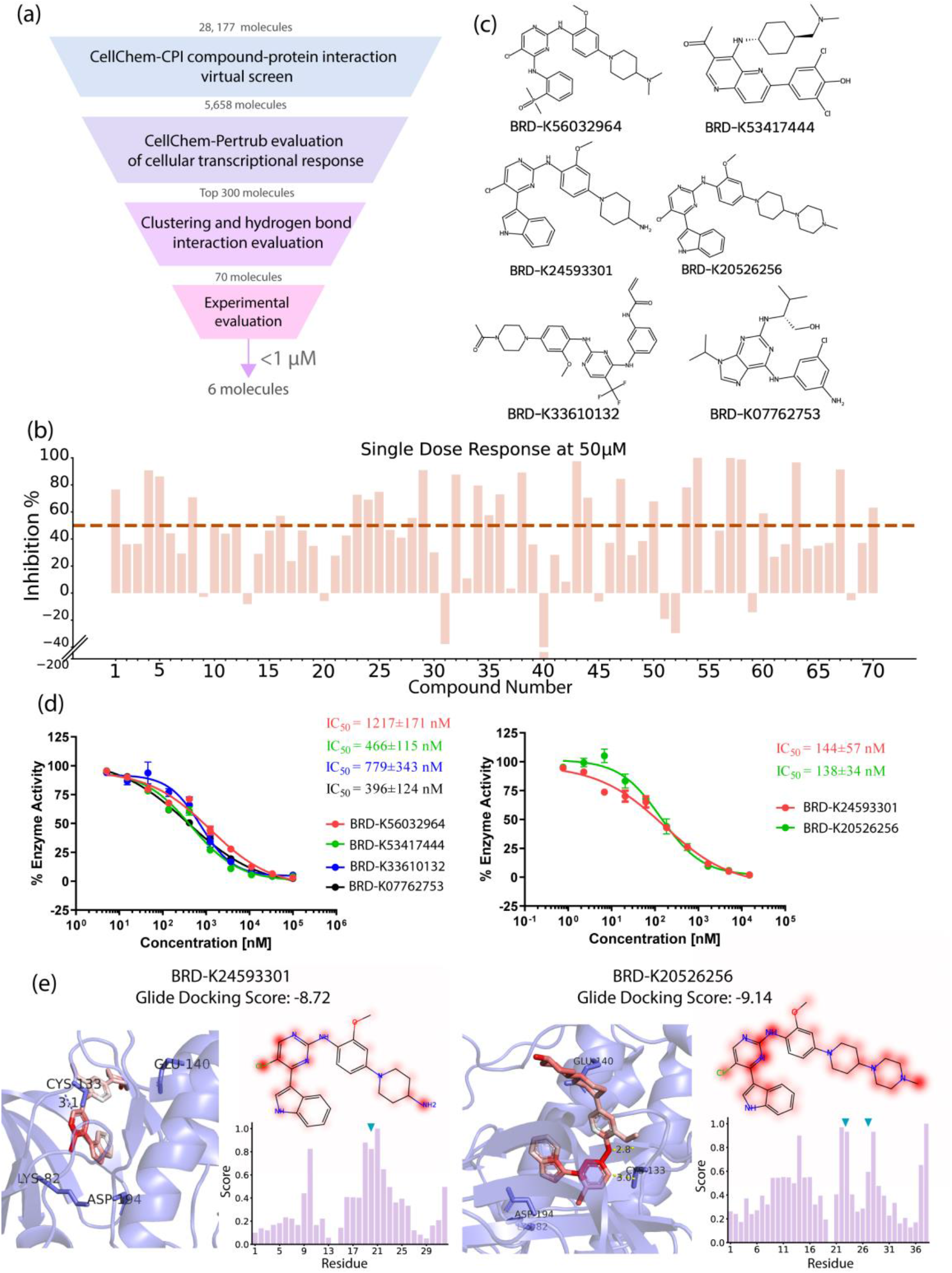
Prospective PLK1 drug repurposing identifies new inhibitors using CellChem molecular representations. (a) The screening workflow illustrates the complete pipeline from computational prediction to experimental validation: 1) PLK1 activity prediction for LINCS database compounds using the CellChem-CPI model. 2) Precise screening of LINCS compounds using the CellChem-Perturb model. 3) Exclusion of known active molecules yielded 70 top-ranked compounds. 4) In vitro enzymatic activity validation system. (b) Chemical structures of six experimentally validated high-activity PLK1 inhibitors (IC_50_ <1 μM). BRD-K24593301 and BRD-K20526256 are highlighted in red boxes (IC_50_ <100 nM). (c) Bar chart of primary screening analysis (50 μM concentration, duplicate testing). The red gradient indicates 28 compounds with >50% inhibition. (d) Dose–response inhibition of PLK1 in an in vitro kinase assay. Percent enzyme activity is plotted against compound concentration (nM, log scale). Points denote measured values (mean ± SD), and solid lines indicate four-parameter logistic fit. (e) Molecular binding pattern. Glide binding mode of BRD-K20526256 with PLK1, highlighting the bidentate hydrogen bond network with CYS133 (blue dashed lines) and the key hydrophobic pocket. Model interpretability validation: the attention-weight heatmap is overlaid on the molecular surface, with red regions indicating automatically recognized pharmacophore features (hydrogen-bond donors/acceptors, marked by triangles in the bar chart).

We selected 9 compounds from these 28 hits for IC_50_ determination. ADP-Glo IC_50_ assays confirmed that 6 of the 9 prioritized compounds had IC_50_ values below 1 μM (Figure 5c). Among these, BRD-K24593301^46, 47^ and BRD-K20526256 were strongly inhibitory(Figure 5d). Although BRD-K20526256 was originally disclosed as an ALK inhibitor, our results suggest that it also exhibits PLK1 inhibitory activity^48^, underscoring the efficacy of CellChem in virtual screening. Consistent with the mechanism-driven organization, the majority of the hits by CellChem were kinase inhibitors. Therefore, molecular embeddings based on transcriptomic responses were enriched for compounds with similar functional profiles. This mechanistic enrichment was further supported by the identification of BRD-K33610132 (Rociletinib)^49^, a clinically advanced phase III EGFR inhibitor, highlighting the potential of CellChem for phenotype-guided drug repurposing.

To assess whether CellChem learns physically meaningful interaction patterns, we compared the interaction sites inferred by CellChem with binding poses generated by Glide docking. The model’s predictions converged on biologically relevant regions of the PLK1 active site. Structural analysis (Figure 5e) indicated that both BRD-K24593301 and BRD-K20526256 engage the functionally critical CYS133 residue in the ATP-binding pocket^50^, confirming that CellChem-prioritized molecules targeted key catalytic features of PLK1.

Consistent with this physical correspondence, the attention-weight visualization (Figure 5e) revealed that the model prioritized pharmacophoric elements, such as hydrogen-bond donors and acceptors that mediate ligand–protein interactions. This alignment between model attention and experimentally supported binding sites demonstrates that CellChem retains mechanistically interpretable interaction logic. Thus, as a cell-guided molecular representation framework, CellChem enables effective screening of active compounds against PLK1 by integrating two complementary levels of evidence: compound–protein interaction prediction (CellChem-CPI) at the molecular binding level and perturbational transcriptomic profile embeddings (CellChem-Perturb) at the cellular response level.

### CellChem efficiently generates perturbation profiles

We developed the CellChem-Generate model, which generates transcriptional profiles for compounds lacking experimentally measured cell response data. These transcriptional profiles can be used to evaluate the effects of compounds on the cell, such as on the transcriptome, and for multi-scale drug discovery that requires cellular response information. Using CellChem molecular representations, we have transferred the jointly contrastive-pretrained chemical-structure encoder and the transcriptional-response encoder to serve as encoders for molecules and cell lines. This allows the model to conditionally generate perturbation transcriptomic profiles in response to compound perturbations (Figure 1d, right, and Supplementary Figure S6). CellChem-Generate was trained on the perturbation dataset curated by Thai-Hoang Pham et al. ^51^ as described in the Methods.

We compared CellChem-Generate with leading models that generate cellular transcriptional profiles using random split, scaffold split, and cell line data-splitting strategies to probe different types of generalization (Figure 6a). Among these, the scaffold split provided the most stringent evaluation, requiring models to extrapolate to chemically novel scaffolds not observed during training. Unlike prior models that rely on non-pretrained or shallow chemical encodings, CellChem uses cell-guided molecular pretraining to learn biologically structured chemical representations for generating transcriptional profiles. Under the scaffold-split setting, CellChem-Generate substantially outperformed all other methods ^51-54^ for generating transcriptional profiles (Figure 6a, middle), indicating that the transcriptome-shaped embedding space supports robust generalization across chemical space rather than mere structural interpolation. The Precision@k analysis ^55^ showed that CellChem-Generate accurately identified the top up- and down-regulated genes, particularly among the top 100 differentially expressed genes (Figure 6b), suggesting better capture of the dominant transcriptional programs induced by chemical perturbations. Full quantitative results for the proposed model and alternative methods are provided in Supplementary Tables 7–9. This ability to generalize is essential for virtual screening, where models must make predictions for novel compounds.

**Figure 6:**
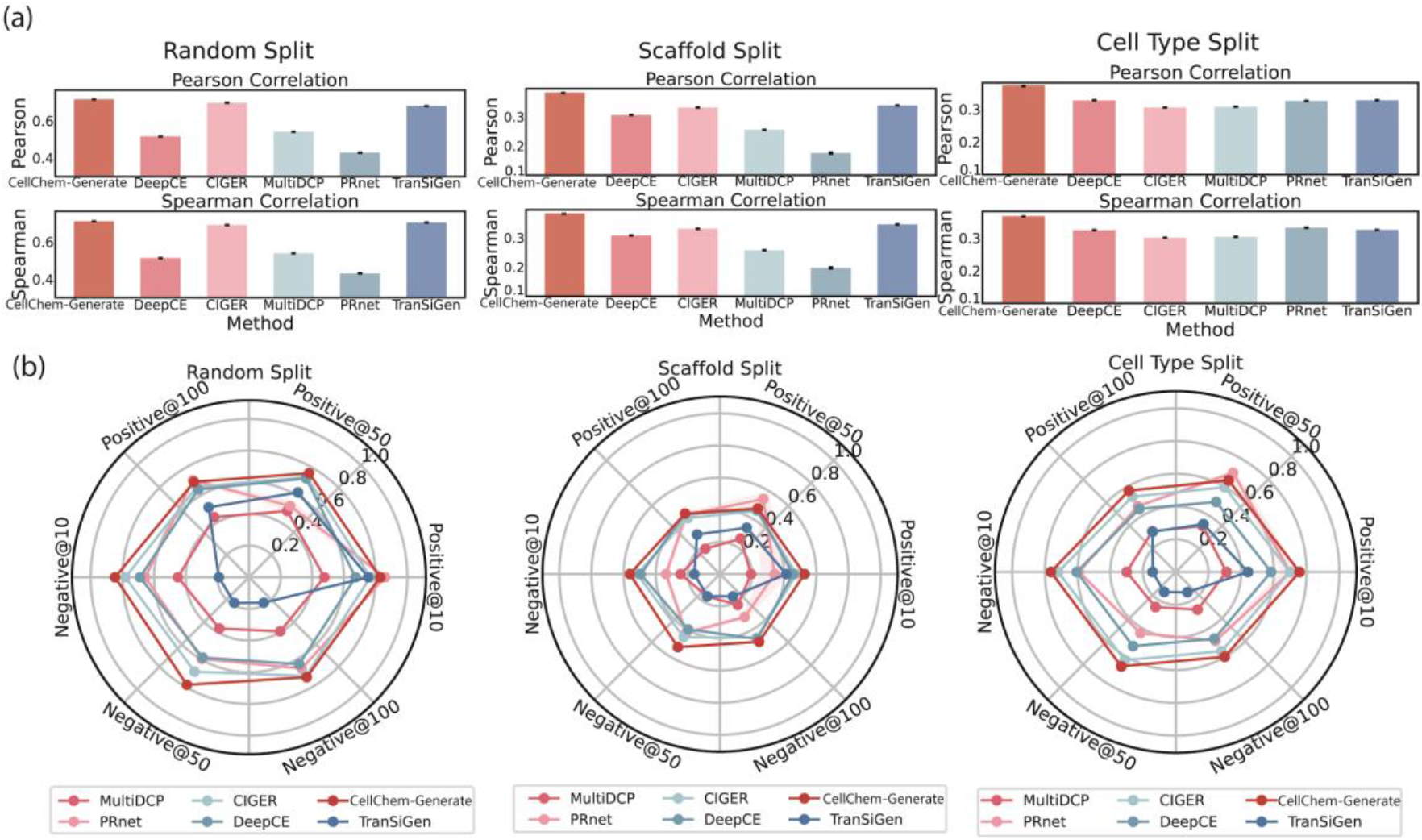
Performance comparison of CellChem-Generate with other models for generating cellular transcriptional profiles. (a) Pearson and Spearman correlations between predicted and actual transcriptional profiles across random, scaffold, and cell-line splits. (b) Negative@precision K and Positive@precision K scores under the three split scenarios.

### GPR6 inverse-agonist drug discovery by CellChem

GPR6 is an orphan GPCR that is expressed in the central nervous system, modulating striatal neural circuits largely through cAMP signaling^56, 57^. Thus, it is a promising therapeutic target for Parkinson’s disease and other neurological disorders. As there are few active chemotypes for GPR6, we aimed to discover inverse agonists with novel chemical scaffolds that are structurally distinct from CVN424, a highly potent GPR6 inverse agonist^58^. GPR6 presents a more challenging screening setting than PLK1 because it lacks reference transcriptional response data from known active compounds, and because GPR6-binding compounds may produce distinct pharmacological effects, including agonism, antagonism, or inverse agonism. We therefore used CellChem-Generate to generate predicted transcriptional profiles and functionally prioritize compounds with likely inverse-agonist-like effects. We conducted a GPR6 inverse-agonist screening using the ChemDiv library. All ChemDiv compounds were represented by CellChem to obtain unified cell-guided molecular embeddings. We used CellChem-CPI to predict CPI scores for compounds and GPR6, and selected compounds with CPI > 0.85, which were then used by CellChem-Generate to predict perturbation-induced transcriptional responses. We compared the responses of these compounds to CVN424 and selected compounds with a perturbation transcriptional response similarity to CVN424 > 0.75. These were tested by molecular docking, drug-likeness filtering^59^, and manual selection, and 51 selected compounds were purchased for the first-round biochemical assay.

The biochemical assay measures a compound’s ability to suppress constitutive GPR6-mediated cAMP signaling. Because it measures changes in basal cAMP rather than responses to an exogenous agonist, it avoids the confounding effects common in antagonist assays, providing a reliable assessment of inverse-agonist activity. Active compounds were defined as those with >50% inhibition of basal cAMP at 50 μM concentration. In the two-dimensional CPI score and perturbation-similarity score plot, the 51 tested compounds are in red in the upper-right corner of the ChemDiv space (Figure 7a). The inhibition by the 51 compounds at 50 μM is illustrated in Figure 7b, with the 14 clustered compounds showing > 50% inhibition. Notably, the 14 experimentally determined compounds show strong enrichment and remain densely clustered within the upper-right subset of these 51 tested compounds (Figure 7c). Combining CellChem-Generate with CellChem-CPI provided strong early enrichment, identifying 8 active compounds within the top 15 ranked compounds. CellChem-Generate alone identified 6/15 and CellChem-CPI alone identified 5/15 compared to the 14/51 that we determined experimentally (Figure 7d). Notably, combining the CellChem-Generate and CellChem-CPI scores was more effective in ranking active compounds than either score alone.

**Figure 7:**
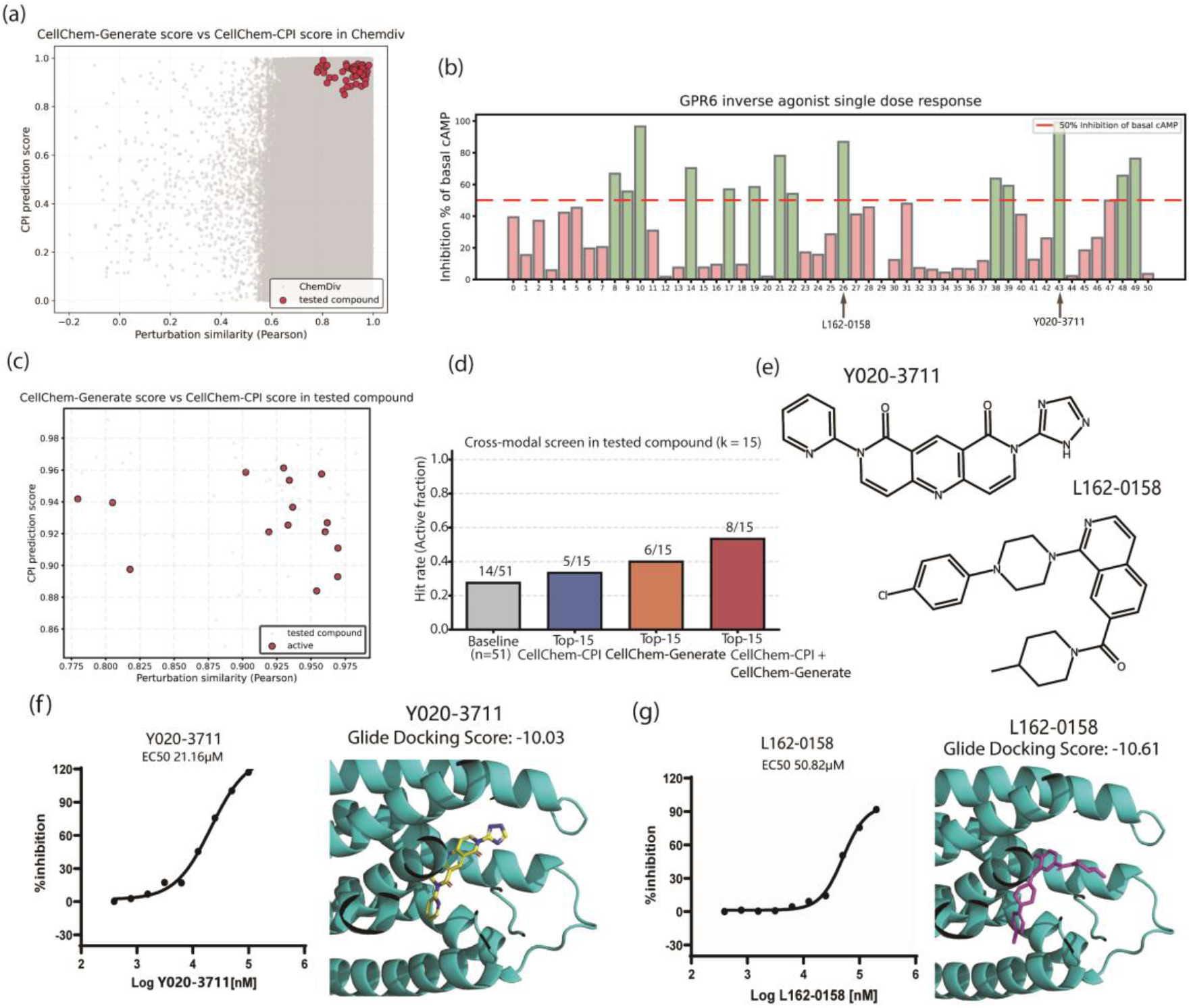
Prospective discovery of GPR6 inverse agonists by CellChem cell-guided molecular representations. (a) Scatter plot of perturbation similarity (Pearson correlation; x-axis) versus CPI prediction score (y-axis) for the ChemDiv compound space (gray). The 51 compounds selected for experimental testing are in red. (b) Biochemical screening assay of the 51 top-ranked compounds at 50 μM, with the dashed line indicating the 50% inhibition threshold used to define active compounds. (c) Same axes as Figure 8a, showing the 51 tested compounds from Figure 8a that were prioritized for experimental testing (red points in Figure 8a). Experimentally confirmed compounds (n = 14; > 50% inhibition) in the cAMP assay are in red. (d) Cross-modal screening improved early enrichment in tested compounds (k = 15). Bar plot showing the hit rate (active fraction) among the 15 top-ranked compounds ranked by CellChem-CPI alone (5/15), CellChem-Generate alone (6/15), or their combined score (8/15). The baseline hit rate across all experimentally tested compounds was 14/51. Combining the two scores yielded the highest early enrichment, indicating improved prioritization of active compounds over either score alone. (e) Chemical structures of representative compounds prioritized by the joint CellChem score (Y020-3711 and L162-0158). (f–g) Dose–response curves for Y020-3711 and L162-0158 confirming inverse agonist activity (EC_50_ in the micromolar range), alongside their Glide docking poses in the GPR6 binding pocket.

We next determined the half-maximal effective concentration (EC_50_) for the six most inhibitory compounds identified by biochemical assays; for compounds Y020-3711 and L162-0158 (Figure 7e), we showed dose-dependent GPR6 inverse agonist activity (Figure 7f, 7g). CVN424, the positive control, had an EC_50_ of 43 nM, similar to the reported value ^60^. Y020-3711 had an EC_50_ of 21.16 μM, and L162-0158 had an EC_50_ of 50.82 μM. Though these two compounds were much less inhibitory than CVN424, they have novel chemical scaffolds with low ECFP4 Tanimoto similarities to CVN424 of 0.192 (L162-0158) and 0.091 (Y020-3711), respectively, and can be optimized as potent GPR6 inverse agonists with novel scaffolds.

### CellChem-Generate predicted cellular transcriptional profiles to map pathways of TCM compounds

Because the molecular targets and pathways affected by traditional Chinese medicine (TCM) compounds are often poorly defined^61-63^, transcriptional profiles can elucidate and categorize the underlying biological mechanisms. Therefore, we used TCM compounds as a validation set for modeling cellular transcriptional responses using CellChem-Generate. TCM therapeutic categories are defined by holistic effects rather than by specific targets or chemical similarity, making them well suited to assess whether CellChem-Generate can produce transcriptomic responses that represent meaningful functional enrichment.

We used interior-warming and heat-clearing medicinal compounds that represent therapeutically antagonistic TCM categories as a case study. The cellular transcriptional profiles of 158 interior-warming TCM compounds and 200 heat-clearing TCM compounds from TCMbank^64^ were generated separately using CellChem-Generate. The predicted transcriptional profiles were embedded into a shared perturbation space and visualized using UMAP^30^ (Figure 8a), and revealed that interior-warming vs. heat-clearing medicinal compounds occupied well-separated regions of the embedding space. This functional separation was quantitatively supported by a linear support vector machine (SVM) classifier trained on the embeddings, which achieved an AUC of 92% in discriminating these two classes (Figure 8a, b). Thus, CellChem-Generate captured clear, clinically relevant distinctions between TCM compounds with opposite therapeutic effects.

**Figure 8:**
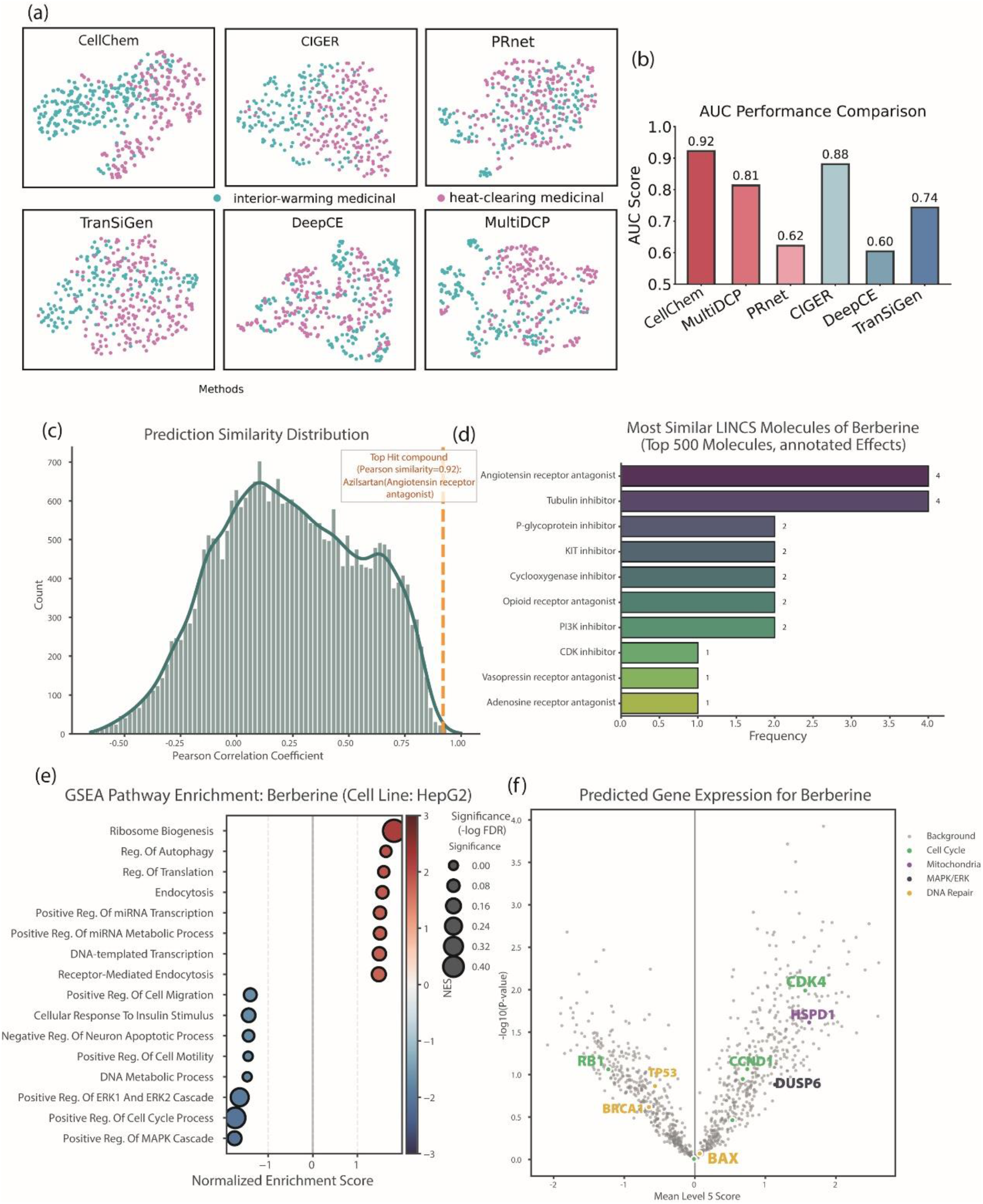
CellChem-Generate reconstructs a system-level berberineprogram in HepG2 cells. (a) Efficacy of various methods in classifying interior-warming and heat-clearing molecules, presented as support vector machine (SVM) AUC scores and a dimensionality reduction visualization. (b) AUC scores for classification of the two therapeutic categories (hemostatic versus blood-activating). (c) Distribution of Pearson correlation coefficients between the CellChem-Generate predicted berberine profile and LINCS reference signatures. The orange dashed line indicates the correlation with the top known hit (an angiotensin receptor antagonist). (d) Top predicted Mechanisms of Action (MOA) categories identified from the nearest neighbors of the predicted berberine signature. (e) Gene Set Enrichment Analysis (GSEA) of the predicted berberine-induced transcriptome in HepG2 cell lines. Bubble size corresponds to the Normalized Enrichment Score (NES), and color intensity indicates statistical significance (-log false discovery rate [FDR]). (f) A volcano plot quantifying gene expression prediction in the HepG2 cell line. The x-axis represents mean prediction intensity, and the y-axis represents consistency.

We then analyzed the HepG2 cell transcriptional profiles for the prototypical alkaloid berberine that were generated by CellChem-Generate. Berberine has anticancer activities, such as the inhibition of proliferation, the promotion of apoptosis, and the induction of cell cycle arrest^65, 66^. It lowers blood pressure by blocking angiotensin receptors, which improves endothelial function and reduces vascular tension ^67^ and has anti-inflammatory activity, including suppression of cyclooxygenase-2 (COX-2) transcription^68, 69^. We calculated the Pearson correlation between the transcriptional profile of berberine generated by CellChem-Generate and reference perturbation profiles from LINCS, and examined the resulting similarity distribution (Figure 8c). The similarity values followed a unimodal distribution centered at about 0.15, indicating that the generated berberine transcriptome occupies a coherent, biologically relevant neighborhood in the LINCS perturbation space. The closest-matching perturbation compound was the angiotensin receptor antagonist azilsartan, with a similarity of 0.92 (Figure 8c). We determined the similarity distribution for the top 500 LINCS reference compounds that remained, after ranking them by their similarity to the berberine-generated transcriptional profile and removing compounds without explicit annotations (Figure 8d). The distribution of similarity scores for compounds across mechanistic categories showed enrichment for cardiovascular regulators (e.g., angiotensin receptor antagonists, vasopressin receptor antagonists), cell cycle/kinase inhibitors (targeting tubulin, KIT, PI3K, and CDK), and inflammatory modulators such as COX inhibitors. These compounds recapitulate the actions of berberine as an angiotensin receptor antagonist (antihypertensive), tubulin/cell cycle inhibitor (antineoplastic), and COX inhibitor (anti-inflammatory).

We then analyzed the predicted effects of berberine in HepG2 cells at the pathway and gene level. Gene Set Enrichment Analysis (GSEA) pathway enrichment revealed downregulation of the MAPK/ERK cascade, cell motility, and cell cycle processes, but upregulation of autophagy pathways (Figure 8e). These predicted pathways align with berberine’s ability to inhibit tumor proliferation and metastasis, induce cell cycle arrest, and trigger autophagy. Analysis of gene transcripts revealed the core regulatory genes (Figure 8f). The predicted upregulation of DUSP6, a negative regulator of ERK signaling, indicated the mechanism for MAPK/ERK pathway suppression^70^. Within the cell cycle and DNA repair modules, increased CDK4 and CCND1 levels coupled with decreased RB1, P53 reflected a disrupted checkpoint network^71^. Thus, the model captured specific cellular responses driving cell cycle arrest rather than general cytotoxic stress. Moreover, the predicted upregulation of mitochondrial chaperone HSPD1 provides target-level evidence for autophagy activation. Overall, the transcriptional profile predicted by CellChem-Generate for berberine perturbation recapitulated its known functions, including antineoplastic activity via cell cycle blockade, antihypertensive effects, and anti-inflammatory regulation, as supported by perturbation-space neighbors, pathway readouts, and gene-regulatory programs.

## Discussion

In this study, we introduce CellChem, a molecular representation learning framework that integrates chemical structures with cellular transcriptional responses. CellChem redefines molecular encoding by embedding both modalities into a shared latent space organized by cellular functions. Unlike conventional representation learning approaches that rely solely on chemical structure, CellChem integrates large-scale transcriptomic response data to reshape the geometry of molecular embeddings, organizing molecules according to functional relationships. Therefore, distances between molecules reflect not only structural similarity but also similarity in the cellular functions they induce. Importantly, this property arises independently of downstream tasks, yielding a molecular space that generalizes across tasks and captures underlying cellular mechanisms. By shifting the representation from chemical space toward cellular space—and from structural similarity toward functional organization—CellChem enables both early enrichment of functionally relevant compounds for CPI prediction and generation of transcriptional response profiles from SMILES that capture pathway- and gene-level effects.

Previous phenotypic screening approaches have incorporated cell-state information into downstream drug discovery, for example, by predicting perturbation signatures from chemical structures or by designing molecules to match transcriptomic profiles^72-74^. However, cellular responses typically enter as downstream conditioning signals: thus, they do not shape the molecular representation space itself. Here, we explicitly combined chemical space with cell response space, using transcriptional responses during pretraining to reshape the geometry of molecular representations that encode both chemical-structure similarity and cellular-functional similarity. Importantly, our model does not simply reconstruct a single perturbation signature but rather uses the predicted transcriptional response as a molecular transcriptional representation that can be reused across tasks. This yields a more universal, task-reusable, and interpretable system for multi-scale drug discovery with cell-guided enhancement. By integrating cross-modal signals in a unified framework, ChemCell outperforms downstream conditioning methods, consistent with our empirical comparisons.

In CellChem, molecular structures, cellular transcriptional profiles, and target-interaction signals are organized within a shared representation space that supports novel drug screening. PLK1 is a well-characterized target with abundant phenotypic and transcriptional evidence, so studying it through structure alone wastes these valuable cellular data. Thus, PLK1 inhibitor discovery is well suited to a multi-scale screening strategy that combines structure-based target engagement with cell-level readouts of functional consequence. For PLK1, CellChem delivers strong early enrichment of functionally relevant kinase inhibitors. In contrast, there is limited data on the transcriptional response to GPR6; however, because cellular guidance distinguishes true pathway suppression from nonspecific binding effects, CellChem is particularly effective for discovering inverse agonists. In the GPR6 study, we used a virtual screening workflow in which CellChem-CPI first identified compounds with binding potential, and CellChem-Generate prioritized them based on predicted transcriptional response patterns consistent with inverse agonism. The GPR6 inverse agonist screening represents a difficult real-world drug discovery scenario. The biology of GPR6 is complex; thus, compounds that can bind GPR6 may be agonists, antagonists, or inverse agonists. Using CellChem-Generate, we can use the predicted cellular transcriptional response data for CVN424 to identify likely inverse agonists. This is a common problem in drug discovery that CellChem-Generate can address by using predicted phenotypes to inform screening. A subset of these compounds demonstrated inverse agonist activity in cellular cAMP assays. In addition, the transcriptional profiles generated by CellChem-Generate provided a functional representation of compounds for drug-discovery applications and insights into their effects on cellular states and pathways. Because CellChem-Generate predicts compound-induced cellular responses at both the gene and pathway levels, it supports function-oriented tasks such as drug repurposing, target prediction, and indication prediction. It is particularly useful for the mechanistic analysis of molecules with unclear targets, such as traditional Chinese medicinal compounds. CellChem-Generate generated transcriptional profiles that organized compounds in therapeutically antagonistic TCM categories into distinct, separable regions in perturbation space. Analysis of the generated transcriptional profile of the prototypical alkaloid berberine supported its described anticancer, antihypertensive, and anti-inflammatory functions.

CellChem’s strength lies in its cell-guided molecular representations, which are organized by biological function. Therefore, it is well suited for applications requiring biological generalization, including off-target effect prediction, polypharmacology, mechanism inference, and phenotypic matching for orphan targets. The current implementation relies on L1000 transcriptomic profiles that include only 978 landmark genes and contain substantial experimental noise, limiting the ability of perturbation profiles to resolve underlying molecular mechanisms. Cell-guided embeddings are also biased toward capturing pathway-level signaling, leading to enrichment of kinase inhibitors among PLK1 hits and a reduced sensitivity to binding modes that do not elicit strong transcriptional responses. For targets lacking perturbation references, such as GPR6, prediction is intrinsically more challenging. Nevertheless, the cross-modal consistency of our results suggests that cell-shaped embeddings by ChemCell exhibit a degree of target specificity beyond their training distribution. Looking ahead, the increasing availability of experimental perturbation profiles generated through RNA-mediated knockdown or CRISPR–Cas9–based gene knockouts of the target gene, followed by transcriptomic profiling, will likely provide direct target-anchored phenotypic constraints. These data should improve the accuracy and reliability of target-focused compound prioritization in this setting.

As high-resolution perturbation atlases and single-cell transcriptomic and proteomic data continue to expand, cell-guided molecular representations are likely to more accurately reflect the true cellular state space. Integrating multi-omics and dynamic perturbation data into CellChem could shift machine learning from static structure-based prediction toward molecular-level simulations of cellular fate. These advances could lead to mechanistically grounded, system-level drug design and the development of computational virtual cell models.

## Methods

### CellChem Training Data Processing

The CMAP database contains four distinct types of gene expression profiles: molecular perturbation profiles, shRNA expression profiles for gene loss-of-function (LoF), wild-type gene cDNA expression profiles for gene overexpression, and CRISPR-based expression profiles for gene loss-of-function. The high-throughput L1000 sequencing measured only 978 landmark genes, representing about 80% of the total gene expression changes in a dataset.

Among the five levels of data processing in the CMAP LINCS 2020 database, we used the most detailed Level 5 data, which contains the post-perturbation expression levels of the 978 genes, their corresponding gene names, perturbation duration, dose concentration, cell line names, and bead batch information at the time of measurement. During data preprocessing, we removed data entries lacking perturbation source molecule information, and we discarded cell lines with too few data points. Our final dataset contained 505,052 perturbation records. For model input, we used the following as condition tokens: cell line, dose concentration, and perturbation duration. Additionally, the bead batch was treated as a batch effect and was removed during training.

### CellChem Architecture

#### Molecular encoder Graph Transformer

In the molecular pretraining module, we adopted a graph transformer architecture, which extends the transformer framework to graph-structured data. Compared with conventional graph attention networks (GATs), graph transformers better preserve and exploit molecular topology when modeling small-molecule graphs. This design enables more effective modeling of long-range dependencies across the molecular graph, thereby strengthening the model’s ability to capture global structural context and, in turn, to represent overall molecular properties.

To effectively capture compound structures and incorporate neighborhood information, we employed the graph transformer to generate node and edge representations for compound graphs. This approach has demonstrated strong performance in graph representation learning. For the embeddings input into the graph transformer, each graph 𝒢 is represented by node features 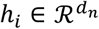 for each node *i* and edge features 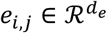 for the connected edge between the node *i* and node *j*. Since the dimensions of node features and edge features differ, we first transformed them into a shared hidden space using a linear transformation, enabling consistent processing in subsequent steps. Specifically,

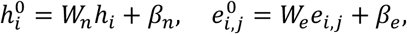

in which 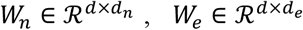 are learnable weight matrices for node and edge representations transformation, respectively. And *β*_*n*_ ∈ ℛ^*d*^ and *β*_*e*_ ∈ ℛ^*d*^ are bias vectors for the linear transformation.

To uncover the spatial neighborhood semantics of each graph and extract fully informative representations for nodes and edges, the embeddings of nodes and edges were iteratively updated using the message-passing paradigm and the multi-head attention mechanism. In the *k*-th single attention head of the *l*-th graph transformer layer, the node and edge embeddings are updated to aggregate the local neighborhood information. Specifically, the attention weight of node *i* with respect to its neighboring node *j* is computed as:

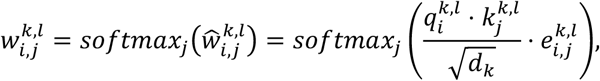

where 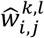 is the attention weight between node *i* and *j* without normalization, and then the query 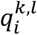, key 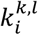, value 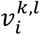 and edge embedding 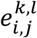 vectors are expressed as,

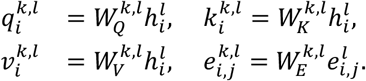

In the above formulas, 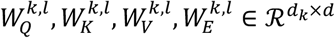 are learnable parameters for the *k*th attention head of the *l*th transformer layer. In particular, 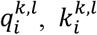 and 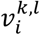 denote query, key, and value vectors for node *i* in the *k*th attention head of the *l*th transformer layer, while 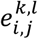 represents the embedding of the connected edge between node *i* and node *j*. Subsequently, we updated node and edge embeddings by incorporating local neighborhood information,

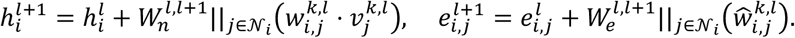

Here, 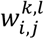 and 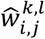 represent the attention weights between node *i* and node *j* in the *k*th head of the *l*th transformer layer, respectively, while 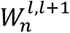 and 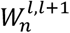 are the learnable weights for node and edge representation propagation in the *l* th layer. Moreover, 𝒩_*i*_ indicates the neighborhood of node *i*, and || represents the concatenation of multiple vectors.

We used a graph transformer architecture with 5 stacked layers, using mean pooling in the pooling layer. The output dimension of the model was set to 512. The batch size was set to 512, and the learning rate was set to 0.005 with a decay of 1e-5, and we employed a contrastive learning loss function.

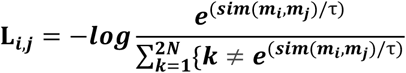

Here τ=0.1, ***m***_***i***_ and ***m***_***j***_ is a vector representation of a pair of positive samples in the hidden space layer. N is the batch size.

#### Cellular transcriptional response encoding (CellChem-Perturb)

For gene expression pretraining on molecular perturbation profiles, we adopted the architecture of a foundation transcriptome model specifically designed for single-cell RNA-sequence analysis that performs strongly across a range of downstream tasks. Here, we extended the input interface by introducing experimental-condition tokens associated with molecular perturbations to better represent the full experimental context. These tokens encode key variables such as perturbation dose, treatment duration, and cell line, enabling the model to learn gene expression changes under diverse and complex experimental conditions. We also explicitly addressed batch effects in RNA expression measurements. In particular, we treated the probe batch used for RNA quantification (the bead batch) as a source of batch effect and included removing this systematic bias as an important training objective.

To encode gene expression values, we discretized expression into 50 bins, thereby reducing noise arising from experimental uncertainty and facilitating stable training. Our implementation was based on scGPT^15^. We modified the tokenization component to support environmental and condition tokens, including perturbation dose, duration, and cell line, allowing the model to capture the influence of these variables and making it well suited for perturbation transcriptomics. To improve optimization stability and generalization, we initialized the model with pretrained parameters from a foundation model pretrained on peripheral blood cells.

#### CellChem for compound-protein interaction prediction models

The positive samples were sourced from the DrugBank data in TransformerCPI^34^. We aligned the DrugBank and CLUE datasets and selected the overlapping data. The number of molecules found was 1062, and the number of proteins was 884. The negative samples were organized as described in HyperAttentionDTI^35^ and shown in detail in Appendix Table 1. For the molecules, the corresponding perturbation data were identified, and for the proteins, the corresponding CRISPR gene knockout data were found. Regarding dataset partitioning strategies, we implemented and analyzed two detailed designs. First, in the initial partitioning method, we randomly assigned one-tenth of the entire dataset as the test set, using the remaining data for five-fold cross-validation. At each fold of the training process, the model generated from that fold was used to predict the test set, and the final evaluation results were computed as the average of the predictions across all folds. In practical applications, predicting interactions between unknown proteins and small molecules is a significant challenge, particularly when high predictive accuracy is required. To address this issue, we designed a second, more complex dataset partitioning strategy. We adopted a scaffold-based partitioning method to ensure the model could handle a more diverse molecular space. Molecules were split into five groups based on their scaffolds, with data from one group used as the test set and the remaining data used for five-fold cross-validation.

Model architecture for the cell response–space CPI model: This model predicts compound– protein interactions directly at the gene expression level by coupling perturbational transcriptomic readouts from compounds with CRISPR-induced transcriptomic readouts from target genes. Following common practice in transcriptomic modeling, landmark-gene expression profiles were discretized into 50 bins to mitigate measurement noise and to better suit Transformer-based encoders. For the compound-side input, when transcriptomic profiles were used as direct inputs, we constructed a compound-level perturbation representation by averaging perturbation profiles across all pert-ids of the same small molecule, aggregating measurements collected across different doses, durations, and cell lines. Both the CRISPR profile and the compound perturbational profile were then embedded and encoded by BERT-style Transformer encoders to produce contextualized sequence embeddings. Finally, a cross-attention block integrated the two modalities to produce an interaction prediction.

Model architecture for the chemical-space CPI model (CellChem-CPI): This model predicts compound–protein interactions from sequence-derived protein representations and structure-derived small-molecule representations, without requiring transcriptomic inputs at inference time. For protein encoding, we used the ESM2 650M protein language model, which captures evolutionary and structural signals from amino-acid sequences and has demonstrated strong performance across diverse protein tasks^75^. For small-molecule encoding, we used CellChem, which outputs a cell-guided molecular representation: transcriptomic information is integrated during CellChem pretraining, enabling inference without requiring cellular transcriptional profiles at test time. After encoding, the compound representation has shape *N* × 512 and the protein representation has shape *L* × 1280. We projected both into a shared 512-dimensional space via linear layers, yielding aligned representations of size *N* × 512 and *L* × 512 for downstream interaction modeling. A cross-attention block then fused the projected protein and compound embeddings to predict the compound–protein interaction.

#### CellChem-Generate Model: a model for generating perturbation profiles from molecular structures

The CellChem generation model is an end-to-end deep learning framework that predicts induced gene perturbation profiles directly from molecular formulas. As illustrated in Figure 1, the overall architecture follows an encoder–decoder paradigm and comprises three core components: a molecular formula encoder, a cell-line encoder, and a decoder, together with the associated training strategy and loss function.

The input to the model is a canonical molecular formula string, and the output is a multi-dimensional vector representing the log-fold-change values for a set of genes, indicating expression changes relative to a control group. To integrate molecular perturbation information into transcriptomic prediction, we designed a cross-modal generation module that conditions gene-level expression modeling on small-molecule embeddings derived from a graph transformer.

Given a gene token sequence **s**, its associated expression values **v**, and a padding mask, transcriptomic representations were first produced using a pretrained cell encoder. Specifically, the encoder maps the input to a sequence of contextualized gene embeddings

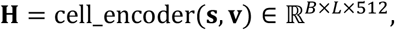

where *B*is the batch size and *L*is the number of gene tokens.

In parallel, each compound was encoded by a graph transformer using its molecular graph *G* = (*X, E*), including atom features, bond types, and optional chemical descriptors. This yielded a compound-level embedding

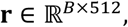

representing the global molecular perturbation state.

To integrate molecular representations with cell-type baseline transcriptomic features, we applied Feature-wise Linear Modulation (FiLM). The molecular embedding **r**was projected to channel-wise scaling and bias vectors,

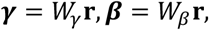

which were broadcast across gene tokens and used to modulate the transcriptomic embeddings:

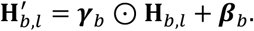

Next, a cross-modal attention mechanism was employed to allow gene tokens to attend to the molecular embedding. Query vectors were obtained from the modulated transcriptomic features, whereas keys and values were derived from the molecular embedding:

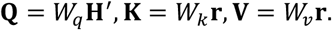

Attention weights were computed as

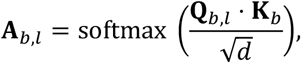

and the attended molecular context was added back to each gene token:

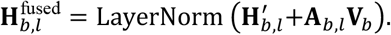

Finally, a gene-wise prediction head was applied to the fused representations to produce a vector of predicted perturbation responses,

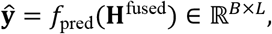

which is aligned with the gene-ordered training targets.

To train the model for accurate prediction of perturbation profiles, the following strategies were employed: masked, centered cosine similarity loss:

We first computed a masked cosine similarity between predicted and observed perturbation vectors. To remove sample-specific expression offsets and focus on relative perturbation patterns, both vectors were mean-centered over valid genes:

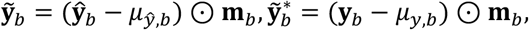

where *μ*_*ŷ,b*_and *μ*_*y,b*_ are the masked means over valid positions.

The cosine similarity for each sample was then computed as

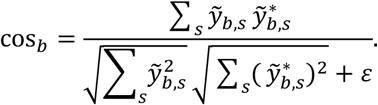

The cosine loss was defined as

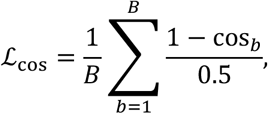

where the scaling factor follows the baseline configuration used in prior work.

Masked Huber (Smooth L1) loss:

To penalize gene-wise deviations while remaining robust to outliers, we additionally applied a masked Huber loss:

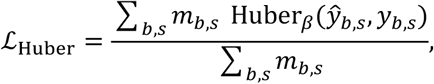

where Huber_*β*_denotes the Smooth L1 loss with transition parameter *β*.

The final training objective was defined as a weighted sum of the two components:

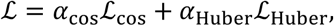

with *α*_cos_ = 1.0and *α*_Huber_ = 0.1.

This composite loss encourages accurate recovery of global perturbation signatures while maintaining stability at the level of individual genes.

The training data for the model were sourced from the dataset curated by Thai-Hoang Pham et al. ^51^ This dataset contains large-scale gene expression profile data from thousands of experiments using various cell lines.

### Evaluation Metrics for Ablation Studies of CellChem-Perturb

#### Clustering separation metrics

The clustering quality was systematically evaluated using three key metrics: the silhouette coefficient (silhouette_score), the Calinski-Harabasz index (calinski_harabasz_score), and the Davies–Bouldin index (davies_bouldin_score). The silhouette coefficient comprehensively measures intra-cluster compactness and inter-cluster separation, with a value range of [-1, 1], where higher values indicate better clustering performance. The Calinski-Harabasz index evaluates clustering quality by computing the ratio of inter-cluster dispersion to intra-cluster dispersion, with higher values representing superior clustering results. The Davies–Bouldin index assesses clustering performance based on the ratio of intra-cluster scatter to inter-cluster distance, with lower values indicating better clustering. These three metrics quantitatively characterize the effects of class separation in the reduced-dimensional space from different perspectives.

#### Intra-Cluster Distance Metrics

Using Euclidean distance (computed via scipy.spatial.distance.pdist) as the measurement standard, we analyzed the distance distribution between all sample pairs within a cluster to derive two key parameters: the average intra-cluster distance (avg_distance), which reflects the overall compactness of the cluster, and the maximum intra-cluster distance (max_distance), which characterizes the separation degree of the most dispersed sample pairs within the cluster. Together, these parameters quantitatively assess the internal structural compactness of the cluster.

#### Inter-Cluster Distance Metrics

Based on the sample data from two clusters (cluster1 and cluster2), we used two calculation methods: maximum distance (max) and average distance (avg). By constructing an inter-sample distance matrix, the separation degree between different clusters was quantified precisely.

This evaluation framework provided a comprehensive quantitative assessment scheme for phenotype-based small molecule models across three dimensions: intra-cluster compactness, inter-cluster separation, and overall clustering performance. All calculations were implemented using standardized algorithms from scipy.spatial.distance, ensuring reliability and reproducibility.

### Methods for Cellular Transcriptional Response Encoding

#### CellChem

This method initially employs the CellChem-Perturb encoder algorithm to encode perturbation profiles, followed by standardized preprocessing of expression data, including log-normalization (sc.pp.log1p) and Z-score standardization (sc.pp.scale). The encoded results were subsequently visualized using the Scanpy module (sc.pl.umap) to generate UMAP dimensionality reduction plots for evaluating encoding efficacy.

#### L1000FWD

The L1000FWD approach employs a cosine-similarity-based weighted graph construction strategy, retaining only the top 0.05% of high-similarity edges, and visualizing the network using the Allegro algorithm. This cosine-similarity-based network construction method comprised four key steps: (1) random sampling of 200,000 sample pairs to compute cosine similarity distributions of feature matrices, with classification-specific connection thresholds established (99.5^th^ percentile for target classification, 90th percentile for cell line classification, and 95^th^ percentile for cmap_name classification); (2) implementation of a streaming batch computation strategy (batch_size=1024) to construct network graphs while retaining only edges exceeding the threshold; (3) removal of sparsely connected nodes through degree filtering; and (4) network visualization using force-directed layout algorithms.

#### SigCom LINCS

This methodology begins by identifying replicate perturbation profiles through metadata matching (including time points, dosage, perturbagens, assay plates, and well IDs), followed by computation of characteristic signatures via the Characteristic Direction (CD) method. The analysis concludes with standardized preprocessing identical to the CellChem approach (log-normalization and Z-score standardization).

Data reprocessing employs the thresholds defined by L1000FWD to filter the dataset, selecting high-quality entries for subsequent clustering analysis.

### Biological Interpretation of Predicted Perturbation Profiles

#### Five-fold cross-validation and ensemble prediction

Predictive performance was evaluated using five-fold cross-validation. The CLUE L1000 Level 5 dataset was partitioned into five subsets, where four were used for training and one for testing in each rotation. During inference, five independent models were loaded to generate transcriptomic perturbation vectors for the target chemical structure. The final prediction was calculated as the mean score of these five outputs to minimize single-model variance and provide a distribution for statistical inference.

#### Pharmacological similarity and functional annotation

Similarity was quantified by calculating Pearson correlation coefficients between the generated expression vectors and the LINCS L1000 reference library. The 500 molecules with the highest correlation values were selected, and entries lacking pharmacological or target annotations were excluded from the analysis. For molecules with multiple annotations, the label strings were split into individual categories. The frequency of each pharmacological class and target function within this selection was then calculated.

#### GSEA pre-ranked enrichment analysis

Gene Set Enrichment Analysis (GSEA) was implemented using the gseapy.prerank algorithm. To prevent ranking ties, a random jitter factor sampled from a normal distribution σ = 10^−6^ was added to the mean scores. Enrichment was tested against the GO_Biological_Process 2023 database, with gene sets limited to sizes between 15 and 1000 genes. Normalized Enrichment Scores (NES) and False Discovery Rates (FDR) were calculated based on 1000 permutations to identify significantly regulated biological processes.

#### Statistical analysis and consistent gene identification

To quantify prediction consistency and filter stochastic noise, a one-sample *t*-test was performed on the five predicted scores for each gene, testing against a null hypothesis of zero. Volcano plots were used to visualize the relationship between effect size (mean score) and statistical significance (− log_10_ *P*-value). Analysis focused on core gene sets associated with the cell cycle, mitochondria, MAPK/ERK, and DNA repair pathways to assess the biological interpretability of the predicted responses.

### Glide docking for PLK1

Molecular docking was performed using the Glide module in standard precision (SP) mode within the Schrödinger software suite. The crystal structure of the human polo-like kinase 1 (PLK1) kinase domain (PDB ID: 2YAC) was retrieved from the Protein Data Bank and employed as the receptor. Protein and ligand preparation were carried out using the Protein Preparation Wizard and LigPrep modules, respectively. The receptor grid was generated using the Glide Grid Generation module and centered on the ATP-binding site of PLK1. Docking was conducted using Glide SP mode with flexible ligand sampling and post-docking minimization. Binding poses were ranked using the GlideScore scoring function, and the top-ranked pose for each ligand was used in the subsequent analysis.

### PLK1 Kinase Inhibition Assay Protocol

This assay determined the half-maximal inhibitory concentration (IC_50_) of test compounds against PLK1 kinase activity, with the known PLK1 inhibitor NMS-1286937 included as a positive control. The assay was performed in triplicate using the ADP-Glo™ luminescence-based kinase detection system in a 96-well low-volume plate.

#### Virtual screening and prioritization pipeline

A LINCS library comprising 28,177 small molecules was first subjected to a GeneLink-based compound–protein interaction (CPI) virtual screening against the target protein, yielding 5,658 compounds with predicted target engagement. These compounds were subsequently evaluated using GeneLink’s perturbation-profile similarity module, in which CellChem-generated transcriptomic signatures were compared against reference phenotypic profiles to quantify functional consistency. The top 300 molecules ranked by the joint CPI and phenotype-similarity score were used for further analysis. To ensure chemical diversity and binding plausibility, these compounds were clustered using molecular fingerprints, followed by analysis of hydrogen-bond interactions within the predicted target-binding pocket, yielding 70 representative compounds. These molecules were then experimentally validated in biochemical assays. Six compounds had activity <1 μM, confirming the effectiveness of the multi-stage GeneLink–CellChem prioritization strategy.

#### Experimental system and conditions

The reaction comprised 1 μL of inhibitor (or 5% dimethyl sulfoxide control), 2 μL of PLK1 enzyme (15 ng/well), and 2 μL of substrate/ATP mixture (final concentrations: 20 μM ATP, 0.2 μg/μL casein). All components were diluted in kinase buffer (40 mM Tris, pH 7.5, 20 mM MgCl_2_, 0.1 mg/mL bovine serum albumin, 50 μM dithiothreitol). After incubation at 25°C for 60 minutes, 5 μL ADP-Glo™ reagent was added, and the reaction was incubated at 25°C for an additional 40 minutes, followed by 10 μL of kinase detection reagent and a further incubation at 25°C for 30 minutes. Luminescence signals were measured with an integration time of 0.5 seconds.

#### Data analysis

Dose–response curves were generated from triplicate experimental data, and the IC_50_ for the compound was calculated to evaluate its inhibition of PLK1 kinase activity.

### Test compound inverse agonist against GPR6-TO by cAMP assay

#### Virtual screening and compound prioritization from a chemical library

To perform large-scale virtual screening against GPR6, we curated the ChemDiv compound library, comprising 1,682,482 commercially available small molecules. All compounds were first embedded using the pretrained CellChem molecular encoder, and their interactions with GPR6 were evaluated using the CellChem-CPI prediction model. The GPR6 transcriptional responses were predicted using the CellChem-Generate model.

The CellChem-CPI model was trained on a GPCR interaction dataset and evaluated using five-fold cross-validation. For each compound–target pair, the predicted interaction scores from the five folds were averaged, and a pair was considered positive if the mean score exceeded 0.85 and the score in every fold was greater than 0.5. In parallel, we identified compounds whose CellChem-Generate–predicted expression profiles were more than 0.75 similar to CVN424. We selected the top 11,646 compounds from this set for subsequent screening.

The prioritized compounds were subjected to structure- and physicochemical-based refinement. We performed ligand preparation with LigPrep to generate appropriate protonation states and conformations, followed by molecular docking with Glide, and retained compounds with docking scores below −9.0. In parallel, we assessed drug-likeness using PCClikeness, choosing compounds with PCClikeness > 0.4. After applying these criteria and choosing only commercially available compounds, we obtained a final set of 51 ChemDiv compounds for experimental validation.

#### Cell Culture and Reagent Preparation

Using the Flp-In-293-GPR6-TO cell line (passage 6), we seeded cells at a density of 2 × 10^6^ cells per flask into 25 cm^2^ flasks and incubated them overnight at 37°C. To induce GPR6 expression, we added 50 ng/mL tetracycline and incubated for a further 3.5 hours. The assay buffer consisted of 1× Hanks’ balanced salt solution, 0.1% fatty acid-free bovine serum albumin, 20 mM HEPES, and 500 μM 3-isobutyl-1-methylxanthine. CVN424, a well-characterized GPR6 inverse agonist^60^, was included as a positive control to calibrate the assay and compare assay performance.

#### Inverse agonist activity assay

Cells were suspended in the assay buffer at a density of 1,000 cells per well, and 15 μL of the cell suspension was added to each well of a 384-well plate, followed by the addition of 5 μL of the test compound, and the plate was incubated at 37°C for 30 minutes. Then, detection reagents were added sequentially: First, the Eu-cAMP tracer was diluted 1:50 in lysis buffer, and 5 μL was added to each well. Next, the Ulight-anti-cAMP antibody was diluted 1:150 with lysis buffer, and 5 μL was added to each well. After this addition, the reaction plate was incubated at room temperature in the dark for 1 hour. Finally, signals were read using an EnVision 2105 plate reader at wavelengths of 615 nm and 665 nm. Inhibition by the compounds was calculated using the following formula: % Inhibition = (Signalcmpd - SignalAve_VC) / (SignalAve_PC - SignalAve_VC) × 100 where Signalcmpd represents the signal of the compound well, SignalAve_VC represents the average signal of the vehicle control wells, and SignalAve_PC represents the average signal of the positive control wells.

## Code availability

The code used for CellChem model training, compound–protein interaction prediction, transcriptional response generation, and downstream analyses is available at https://github.com/Chenjxjx/CellChem.

## Acknowledgements

This work was supported in part by the Innovative Drug Research and Development National Science and Technology Major Project (No. 2025ZD1803103), the National Natural Science Foundation of China (Grant Nos. 220330010 and T2321001), the Major Project of Guangzhou National Laboratory (Grant No. GZNL2024A01005), the CAMS Innovation Fund for Medical Sciences (2021-I2M-5-014), and the Anhui’s Plans for Major Provincial Science Technology Projects (202303a07020009). Part of the analysis was performed on the High Performance Computing Platform of the Center for Life Sciences, Peking University.

## References

1. Wang, Y., Wang, J., Cao, Z. & Barati Farimani, A. Molecular contrastive learning of representations via graph neural networks. Nature Machine Intelligence 4, 279–287 (2022).

2. Wang, X. et al. Multimodal pre-training models of molecular representation for drug discovery. National Science Review 13, nwaf495 (2026).

3. Zhang, P. et al. A deep learning framework for in silico screening of anticancer drugs at the single-cell level. National science review 12, nwae451 (2025).

4. Guo, F. et al. Foundation models in bioinformatics. National science review 12, nwaf028 (2025).

5. Liu, S. et al. Pre-training molecular graph representation with 3d geometry. arXiv preprint arXiv:2110.07728 (2021).

6. Fang, Y. et al. Knowledge graph-enhanced molecular contrastive learning with functional prompt. Nature Machine Intelligence 5, 542–553 (2023).

7. Subramanian, A. et al. A next generation connectivity map: L1000 platform and the first 1,000,000 profiles. Cell 171, 1437–1452. e1417 (2017).

8. Behar, M., Barken, D., Werner, S.L. & Hoffmann, A. The dynamics of signaling as a pharmacological target. Cell 155, 448–461 (2013).

9. Meng, J., Lu, M., Chen, Y., Gao, S.-J. & Huang, Y. Robust inference of the context specific structure and temporal dynamics of gene regulatory network. BMC genomics 11, S11 (2010).

10. Kana, O. et al. Generative modeling of single-cell gene expression for dose-dependent chemical perturbations. Patterns 4 (2023).

11. Van Tilborg, D., Alenicheva, A. & Grisoni, F. Exposing the limitations of molecular machine learning with activity cliffs. Journal of chemical information and modeling 62, 5938–5951 (2022).

12. Stumpfe, D., Hu, H. & Bajorath, J. Evolving concept of activity cliffs. ACS omega 4, 14360–14368 (2019).

13. Bunne, C. et al. How to build the virtual cell with artificial intelligence: Priorities and opportunities. Cell 187, 7045–7063 (2024).

14. Tang, L. The virtual cell. Nature Methods 22, 2493–2493 (2025).

15. Cui, H. et al. scGPT: toward building a foundation model for single-cell multi-omics using generative AI. Nature Methods, 1–11 (2024).

16. Theodoris, C.V. et al. Transfer learning enables predictions in network biology. Nature 618, 616–624 (2023).

17. Hao, M. et al. Large-scale foundation model on single-cell transcriptomics. Nature methods 21, 1481–1491 (2024).

18. Tong, X. et al. Deep representation learning of chemical-induced transcriptional profile for phenotype-based drug discovery. Nature Communications 15, 5378 (2024).

19. Cheng, J. et al. GexMolGen: cross-modal generation of hit-like molecules via large language model encoding of gene expression signatures. Briefings in Bioinformatics 25, bbae525 (2024).

20. Méndez-Lucio, O., Baillif, B., Clevert, D.-A., Rouquié, D. & Wichard, J. De novo generation of hit-like molecules from gene expression signatures using artificial intelligence. Nature communications 11, 10 (2020).

21. Kim, H. et al. A genotype-to-drug diffusion model for generation of tailored anti-cancer small molecules. Nature Communications 16, 5628 (2025).

22. Zhu, J. et al. Prediction of drug efficacy from transcriptional profiles with deep learning. Nature biotechnology 39, 1444–1452 (2021).

23. DeMeo, B. et al. Active learning framework leveraging transcriptomics identifies modulators of disease phenotypes. Science 390, eadi8577 (2025).

24. Tian, Y. et al. What makes for good views for contrastive learning? Advances in neural information processing systems 33, 6827–6839 (2020).

25. Yun, S., Jeong, M., Kim, R., Kang, J. & Kim, H.J. Graph transformer networks. Advances in neural information processing systems 32 (2019).

26. Wang, Z., Lachmann, A., Keenan, A.B. & Ma’Ayan, A. L1000FWD: fireworks visualization of drug-induced transcriptomic signatures. Bioinformatics 34, 2150–2152 (2018).

27. Evangelista, J.E. et al. SigCom LINCS: data and metadata search engine for a million gene expression signatures. Nucleic acids research 50, W697–W709 (2022).

28. Barrett, S.D. et al. The discovery of the benzhydroxamate MEK inhibitors CI-1040 and PD 0325901. Bioorganic & medicinal chemistry letters 18, 6501–6504 (2008).

29. Erlichman, C. Tanespimycin: the opportunities and challenges of targeting heat shock protein 90. Expert opinion on investigational drugs 18, 861–868 (2009).

30. McInnes, L., Healy, J. & Melville, J. Umap: Uniform manifold approximation and projection for dimension reduction. arXiv preprint arXiv:1802.03426 (2018).

31. Löwe, J. et al. Crystal structure of the 20 S proteasome from the archaeon T. acidophilum at 3.4 Å resolution. Science 268, 533–539 (1995).

32. Hubbard, S.R., Wei, L. & Hendrickson, W.A. Crystal structure of the tyrosine kinase domain of the human insulin receptor. Nature 372, 746–754 (1994).

33. Hadipour, H. et al. GraphBAN: An inductive graph-based approach for enhanced prediction of compound-protein interactions. Nature Communications 16, 2541 (2025).

34. Chen, L. et al. TransformerCPI: improving compound–protein interaction prediction by sequence-based deep learning with self-attention mechanism and label reversal experiments. Bioinformatics 36, 4406–4414 (2020).

35. Zhao, Q., Zhao, H., Zheng, K. & Wang, J. HyperAttentionDTI: improving drug– protein interaction prediction by sequence-based deep learning with attention mechanism. Bioinformatics 38, 655–662 (2022).

36. Nguyen, T. et al. GraphDTA: predicting drug–target binding affinity with graph neural networks. Bioinformatics 37, 1140–1147 (2021).

37. Huang, K., Xiao, C., Glass, L.M. & Sun, J. MolTrans: molecular interaction transformer for drug–target interaction prediction. Bioinformatics 37, 830–836 (2021).

38. Knox, C. et al. DrugBank 6.0: the DrugBank knowledgebase for 2024. Nucleic acids research 52, D1265–D1275 (2024).

39. Hofheinz, R.-D. et al. An open-label, phase I study of the polo-like kinase-1 inhibitor, BI 2536, in patients with advanced solid tumors. Clinical Cancer Research 16, 4666–4674 (2010).

40. Frost, A. et al. Phase i study of the Plk1 inhibitor BI 2536 administered intravenously on three consecutive days in advanced solid tumours. Current oncology 19, e28 (2012).

41. Sebastian, M. et al. The efficacy and safety of BI 2536, a novel Plk-1 inhibitor, in patients with stage IIIB/IV non-small cell lung cancer who had relapsed after, or failed, chemotherapy: results from an open-label, randomized phase II clinical trial. Journal of Thoracic Oncology 5, 1060–1067 (2010).

42. Friesner, R.A. et al. Glide: a new approach for rapid, accurate docking and scoring. 1. Method and assessment of docking accuracy. Journal of medicinal chemistry 47, 1739–1749 (2004).

43. Friesner, R.A. et al. Extra precision glide: Docking and scoring incorporating a model of hydrophobic enclosure for protein™ ligand complexes. Journal of medicinal chemistry 49, 6177–6196 (2006).

44. Gaulton, A. et al. ChEMBL: a large-scale bioactivity database for drug discovery. Nucleic acids research 40, D1100–D1107 (2012).

45. Zhu, T. et al. Hit identification and optimization in virtual screening: practical recommendations based on a critical literature analysis. J Med Chem 56, 6560–6572 (2013).

46. Wang, Y. et al. Novel ALK inhibitor AZD3463 inhibits neuroblastoma growth by overcoming crizotinib resistance and inducing apoptosis. Scientific reports 6, 19423 (2016).

47. Moharram, S.A., Shah, K., Khanum, F., Rönnstrand, L. & Kazi, J.U. The ALK inhibitor AZD3463 effectively inhibits growth of sorafenib-resistant acute myeloid leukemia. Blood cancer journal 9, 5 (2019).

48. Aquila, B. et al. (World Intellectual Property Organization, Geneva; 2012).

49. Sequist, L.V. et al. Rociletinib in EGFR-mutated non–small-cell lung cancer. New England Journal of Medicine 372, 1700–1709 (2015).

50. Xie, W. et al. Accelerating discovery of bioactive ligands with pharmacophore-informed generative models. Nature communications 16, 2391 (2025).

51. Pham, T.-H. et al. Chemical-induced gene expression ranking and its application to pancreatic cancer drug repurposing. Patterns 3 (2022).

52. Wu, Y., Liu, Q., Qiu, Y. & Xie, L. Deep learning prediction of chemical-induced dose-dependent and context-specific multiplex phenotype responses and its application to personalized alzheimer’s disease drug repurposing. PLoS computational biology 18, e1010367 (2022).

53. Pham, T.-H., Qiu, Y., Zeng, J., Xie, L. & Zhang, P. A deep learning framework for high-throughput mechanism-driven phenotype compound screening and its application to COVID-19 drug repurposing. Nature machine intelligence 3, 247–257 (2021).

54. Qi, X. et al. Predicting transcriptional responses to novel chemical perturbations using deep generative model for drug discovery. Nature Communications 15, 9256 (2024).

55. Kar, P., Narasimhan, H. & Jain, P. in International Conference on Machine Learning 189–198 (PMLR, 2015).

56. Shrader, S.H. & Song, Z.H. Discovery of endogenous inverse agonists for G protein-coupled receptor 6. Biochem Biophys Res Commun 522, 1041–1045 (2020).

57. Sun, H. et al. First-Time Disclosure of CVN424, a Potent and Selective GPR6 Inverse Agonist for the Treatment of Parkinson’s Disease: Discovery, Pharmacological Validation, and Identification of a Clinical Candidate. Journal of Medicinal Chemistry 64, 9875–9890 (2021).

58. Brice, N.L. et al. Development of CVN424: A Selective and Novel GPR6 Inverse Agonist Effective in Models of Parkinson Disease. J Pharmacol Exp Ther 377, 407–416 (2021).

59. Bickerton, G.R., Paolini, G.V., Besnard, J., Muresan, S. & Hopkins, A.L. Quantifying the chemical beauty of drugs. Nature Chemistry 4, 90–98 (2012).

60. Sun, H. et al. First-time disclosure of CVN424, a potent and selective GPR6 inverse agonist for the treatment of Parkinson’s disease: discovery, pharmacological validation, and identification of a clinical candidate. Journal of Medicinal Chemistry 64, 9875–9890 (2021).

61. Yoo, M. et al. Exploring the molecular mechanisms of Traditional Chinese Medicine components using gene expression signatures and connectivity map. Computer methods and programs in biomedicine 174, 33–40 (2019).

62. Yu-Ping, T. et al. Modern research thoughts and methods on bio-active components of TCM formulae. Chinese Journal of Natural Medicines 20, 481–493 (2022).

63. Gu, X., Hao, D. & Xiao, P. Research progress of Chinese herbal medicine compounds and their bioactivities: Fruitful 2020. Chinese herbal medicines 14, 171–186 (2022).

64. Lv, Q. et al. TCMBank: bridges between the largest herbal medicines, chemical ingredients, target proteins, and associated diseases with intelligence text mining. Chemical science 14, 10684–10701 (2023).

65. Wang, Y., Liu, Y., Du, X., Ma, H. & Yao, J. The Anti-Cancer Mechanisms of Berberine: A Review. Cancer Manag Res 12, 695–702 (2020).

66. Qi, H.W., Xin, L.Y., Xu, X., Ji, X.X. & Fan, L.H. Epithelial-to-mesenchymal transition markers to predict response of Berberine in suppressing lung cancer invasion and metastasis. J Transl Med 12, 22 (2014).

67. Feng, X. et al. Berberine in Cardiovascular and Metabolic Diseases: From Mechanisms to Therapeutics. Theranostics 9, 1923–1951 (2019).

68. Fukuda, K. et al. Inhibition by berberine of cyclooxygenase-2 transcriptional activity in human colon cancer cells. J Ethnopharmacol 66, 227–233 (1999).

69. Kuo, C.L., Chi, C.W. & Liu, T.Y. The anti-inflammatory potential of berberine in vitro and in vivo. Cancer Lett 203, 127–137 (2004).

70. Liang, K.W. et al. Berberine suppresses MEK/ERK-dependent Egr-1 signaling pathway and inhibits vascular smooth muscle cell regrowth after in vitro mechanical injury. Biochem Pharmacol 71, 806–817 (2006).

71. Liu, Z. et al. Berberine induces p53-dependent cell cycle arrest and apoptosis of human osteosarcoma cells by inflicting DNA damage. Mutat Res 662, 75–83 (2009).

72. Guo, Y. et al. Modelling drug-induced cellular perturbation responses with a biologically informed dual-branch transformer. Nature Machine Intelligence, 1–17 (2026).

73. Xing, J. et al. Deep-learning-based de novo discovery and design of therapeutics that reverse disease-associated transcriptional phenotypes. Cell (2026).

74. DeMeo, B. et al. Active learning framework leveraging transcriptomics identifies modulators of disease phenotypes. Science 390, eadi8577 (2025).

75. Lin, Z. et al. Evolutionary-scale prediction of atomic-level protein structure with a language model. Science 379, 1123–1130 (2023).

