## Supplemental information for "CellChem: Cellular transcriptional responses reshape molecular representation space for efficient and multi-scale drug discovery"

**Supplementary Figures**

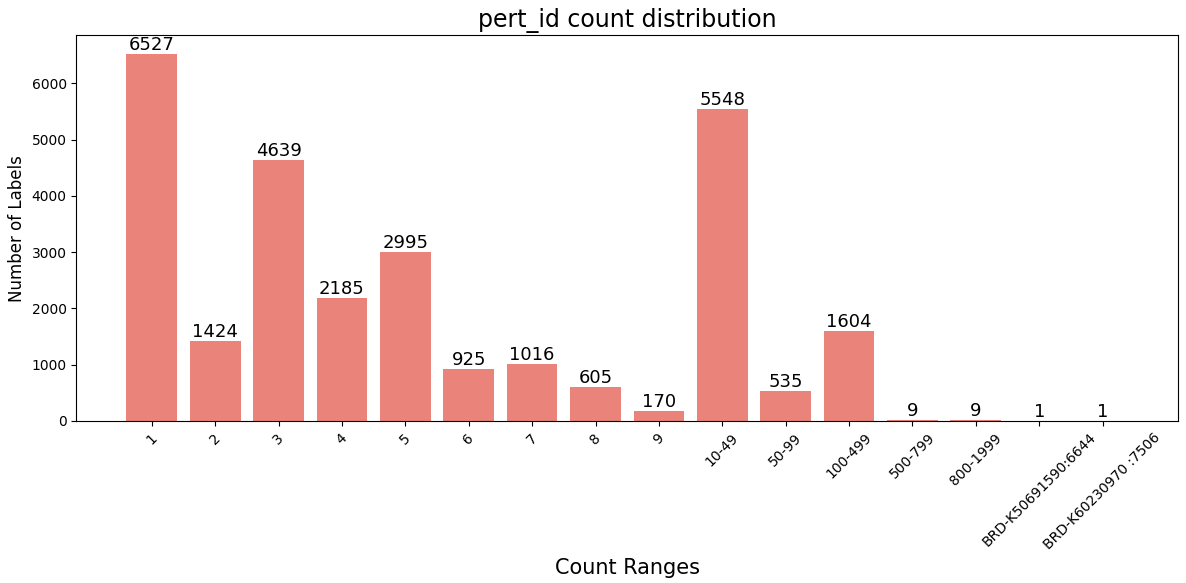

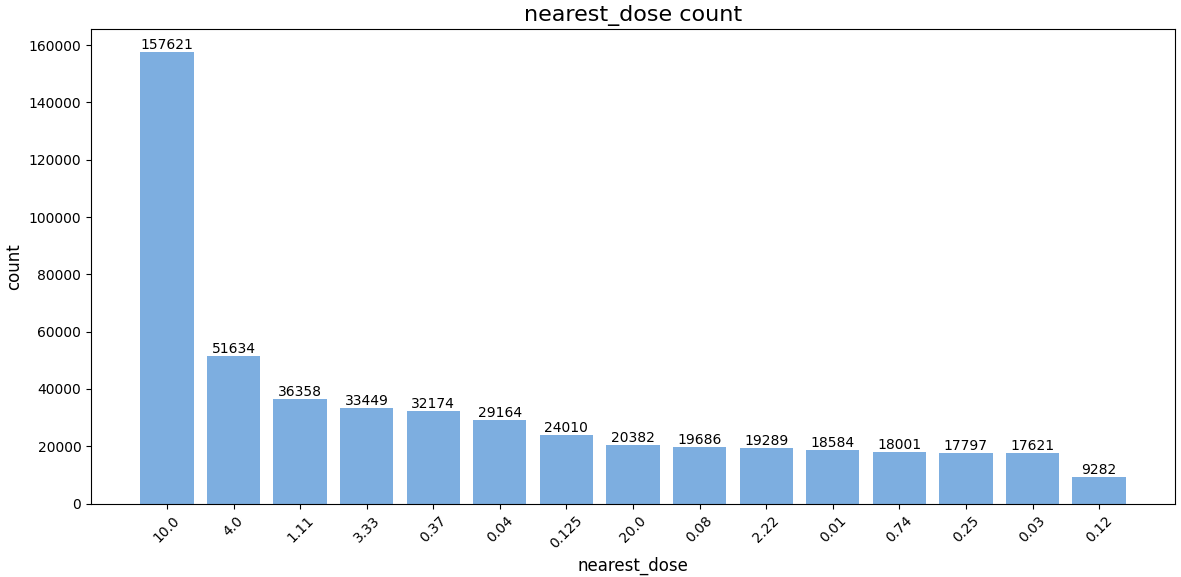

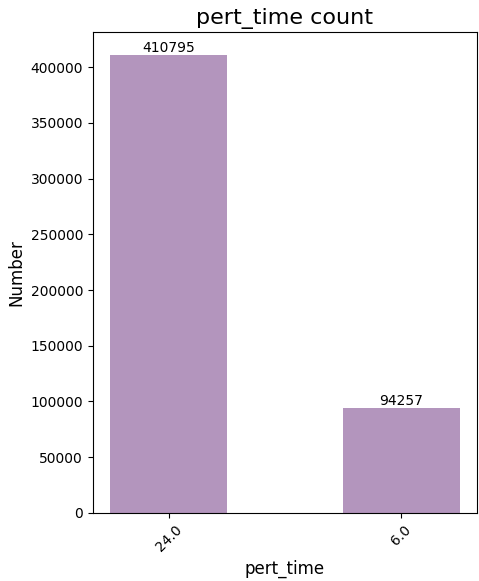

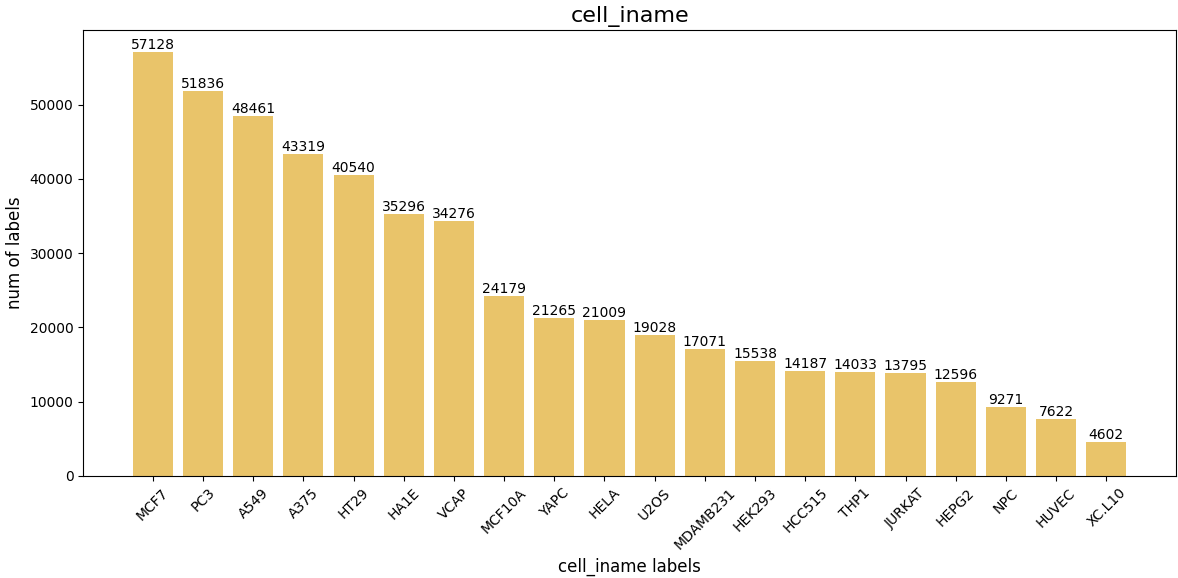

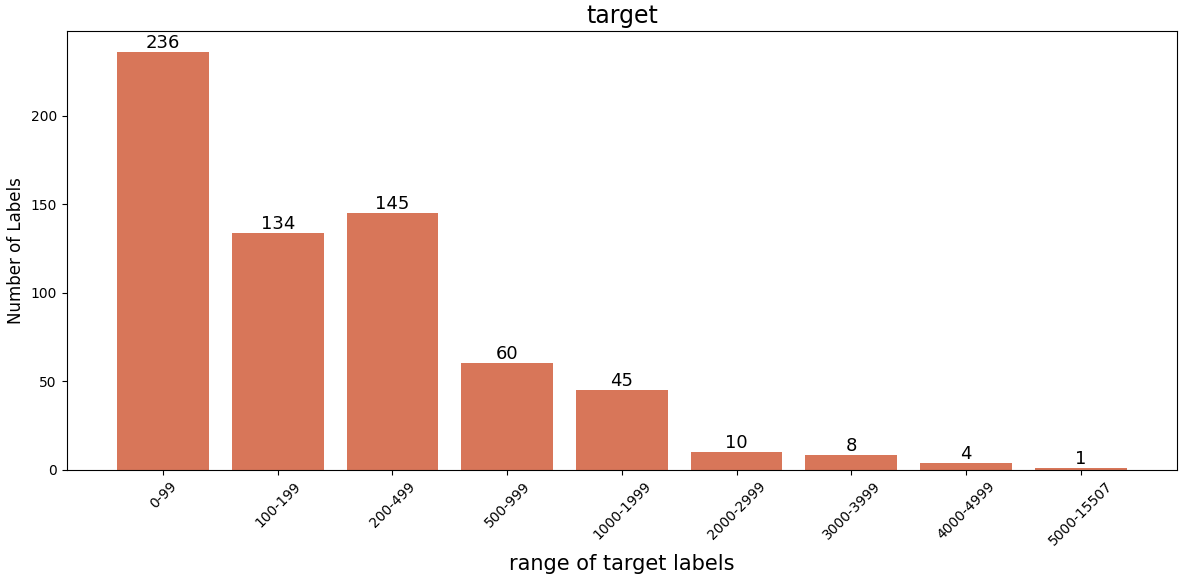

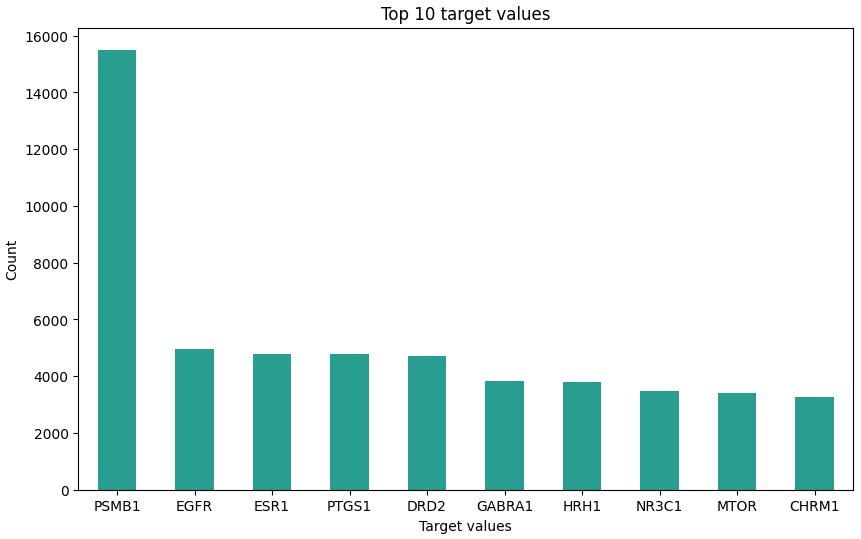

**Figure 1: The molecule perturbation dataset used during pretraining includes statistical results based on labels such as ID, dose, perturbation duration, and cell line.**

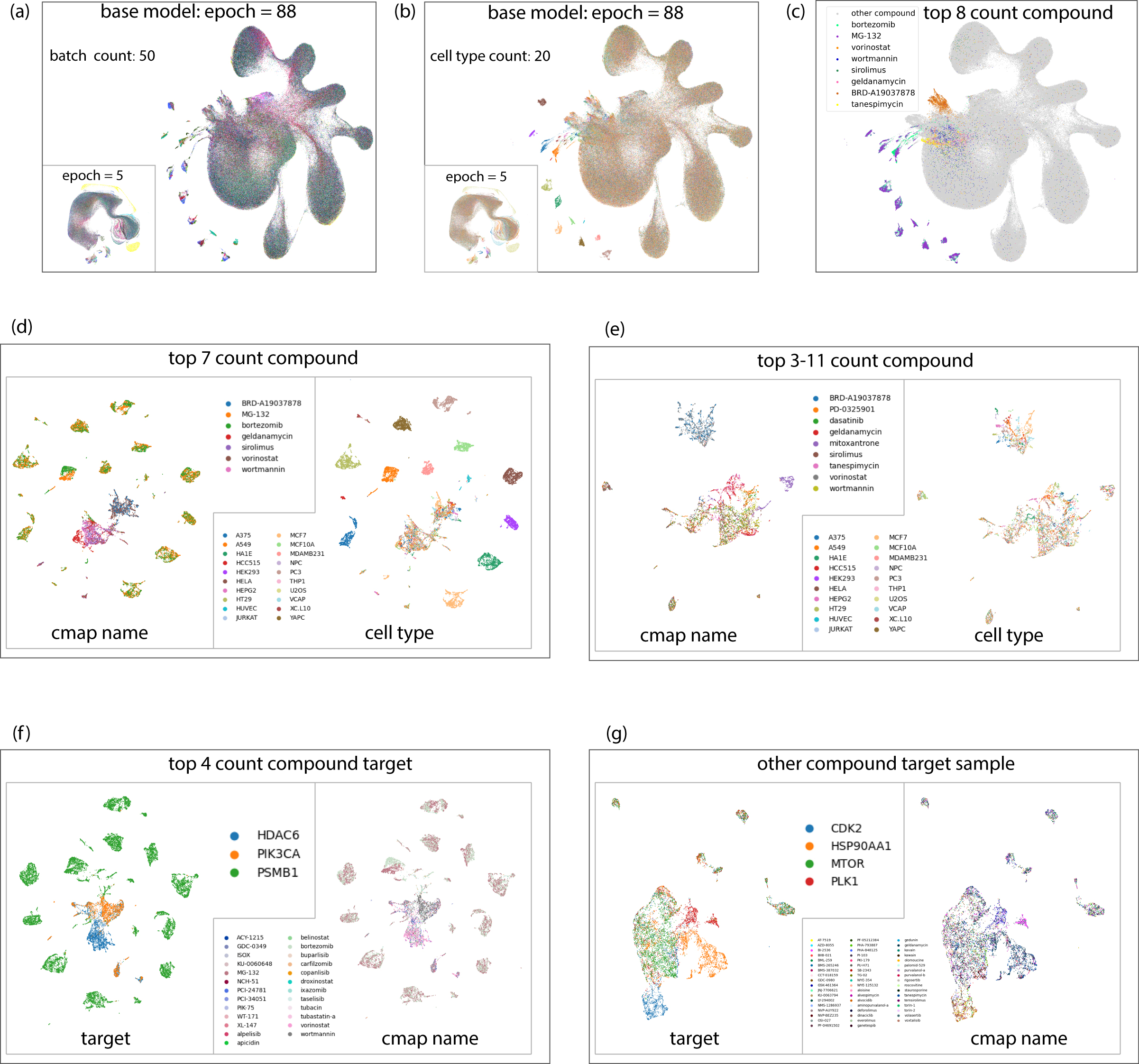

**Figure 2: Dimensionality reduction clustering results for gene expression profiles, fine-tuned by the model on different data types.** (a) The subfigure shows the dimensionality reduction results at the 5th epoch, and the large figure shows the results at the 88th epoch. The colors represent different data batches. (b) The subfigure shows the dimensionality reduction results at the 5th epoch, and the large figure shows the results at the 88th epoch. The colors represent different cell types. The colors distinguish between the top 8 small molecules with the most perturbation expression profile data. (c) The left image shows dimensionality reduction clustering of the top 7 small molecules with the most perturbation expression profiles, colored by molecular type. The right image shows the same clustering but colored by cell type. (d) The left image shows dimensionality reduction clustering of small molecules ranked 3rd to 11th in terms of perturbation expression profile data, colored by molecular type. The right image shows the same clustering but colored by cell type. (e) The left image shows dimensionality reduction clustering of the top 3 targets with the most perturbation expression profile data, colored by target type. The right image shows the same clustering but colored by molecular type. (f) Dimensionality reduction clustering for other targets (except for panel C), with the left image colored by target type and the right image colored by molecular type.

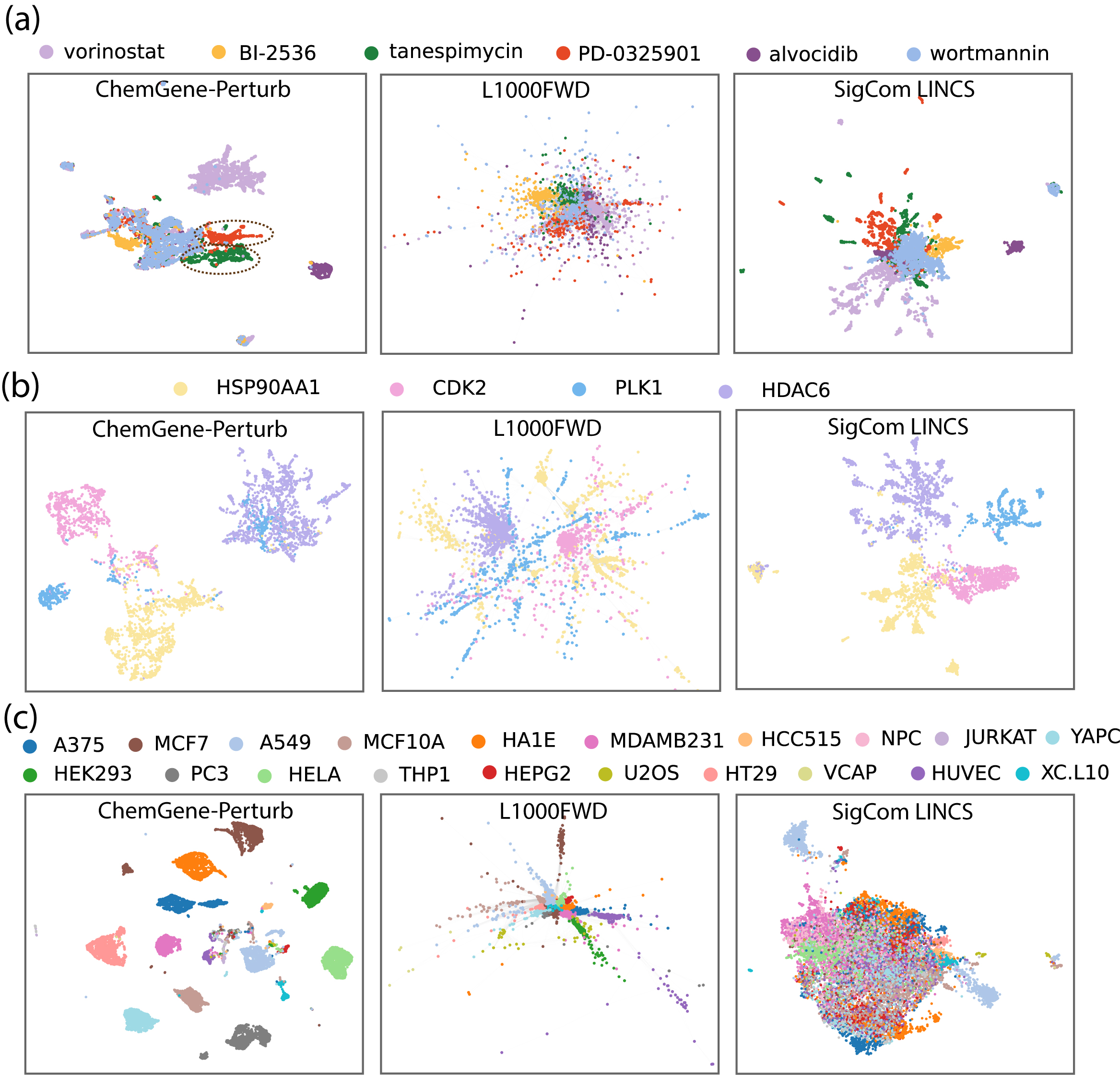

**Figure 3：Comparative Analysis of CellChem and Other Perturbation Profiling Embedding Methods via UMAP Visualization across Molecular, Target, and Cellular Dimensions.** (a) Molecular level, (b) target level, (c) cell-line level.

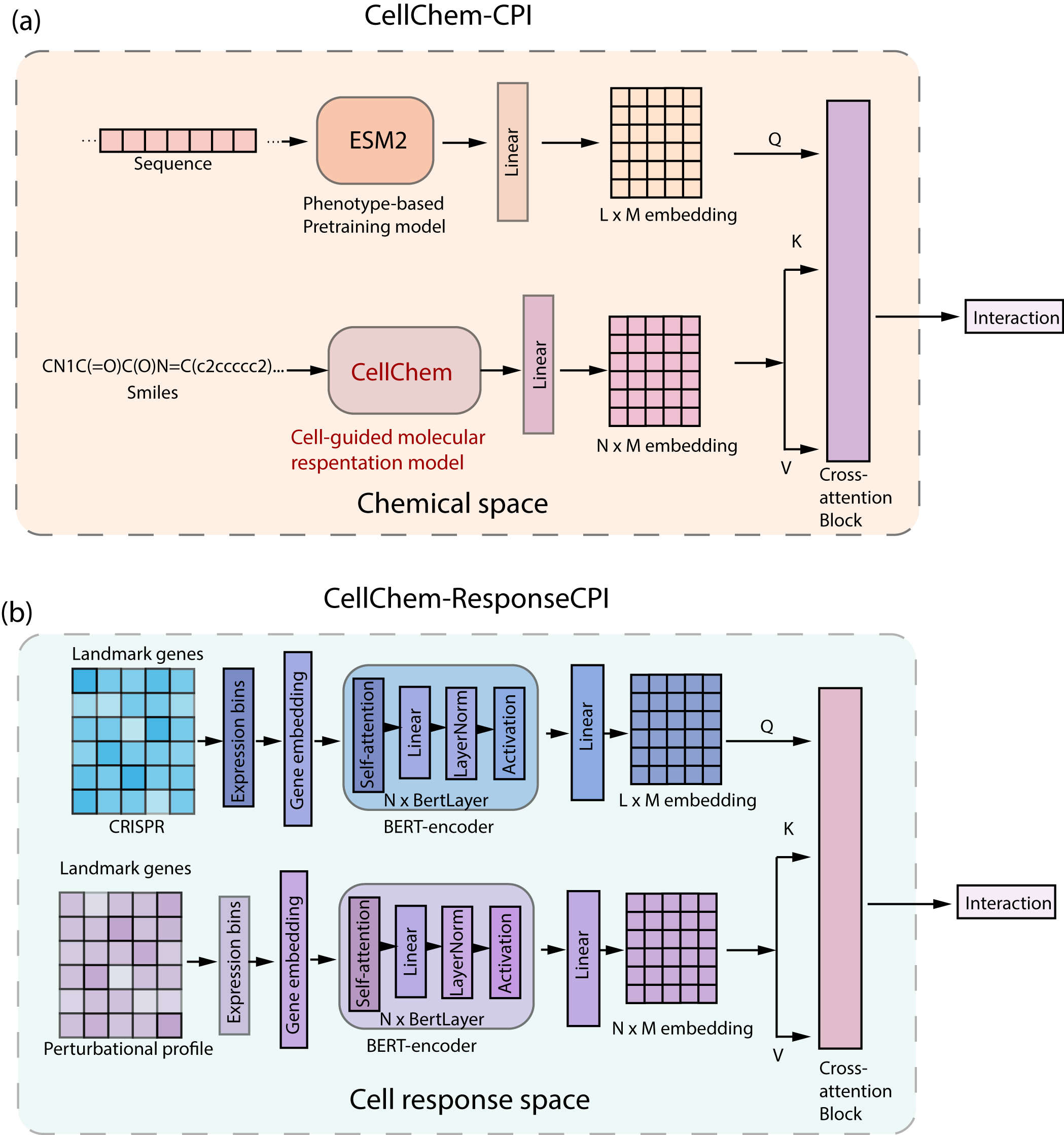

**Figure 4: CellChem-CPI's architecture and CellChem-ResponseCPI.** (a) CellChem-CPI. The CPI model takes protein sequence and cell-guided CellChem molecular representation as inputs. Protein sequences are encoded by Evolutionary Scale Modeling (ESM). The resulting embeddings are projected and integrated via the same cross-attention block to predict compound–protein interaction. Transcriptomic information is integrated during CellChem pretraining, enabling inference without requiring cellular transcriptional profiles at test time. (b) CellChem-ResponseCPI. The CPI model takes transcriptional profiles as inputs: a CRISPR perturbation profile as a protein-side readout and a compound-induced perturbational transcriptomic profile as a compound-side readout. Each profile is discretized into expression bins, embedded, and encoded by BERT-style encoders to produce sequence-level embeddings, which are then fused by a cross-attention block to predict compound–protein interaction.

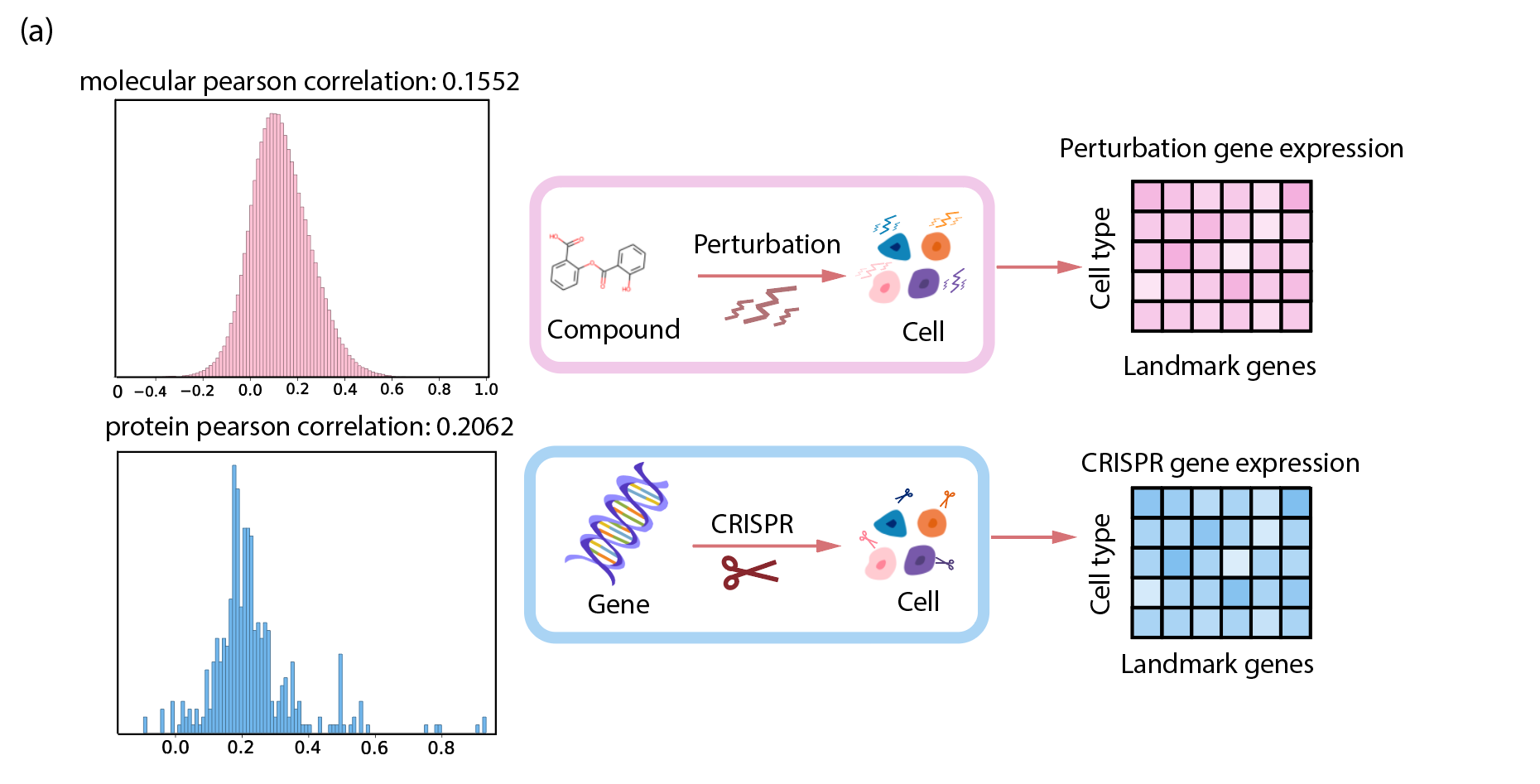

**Figure 5: The molecule perturbation dataset used during pretraining includes statistical results based on labels such as target.**

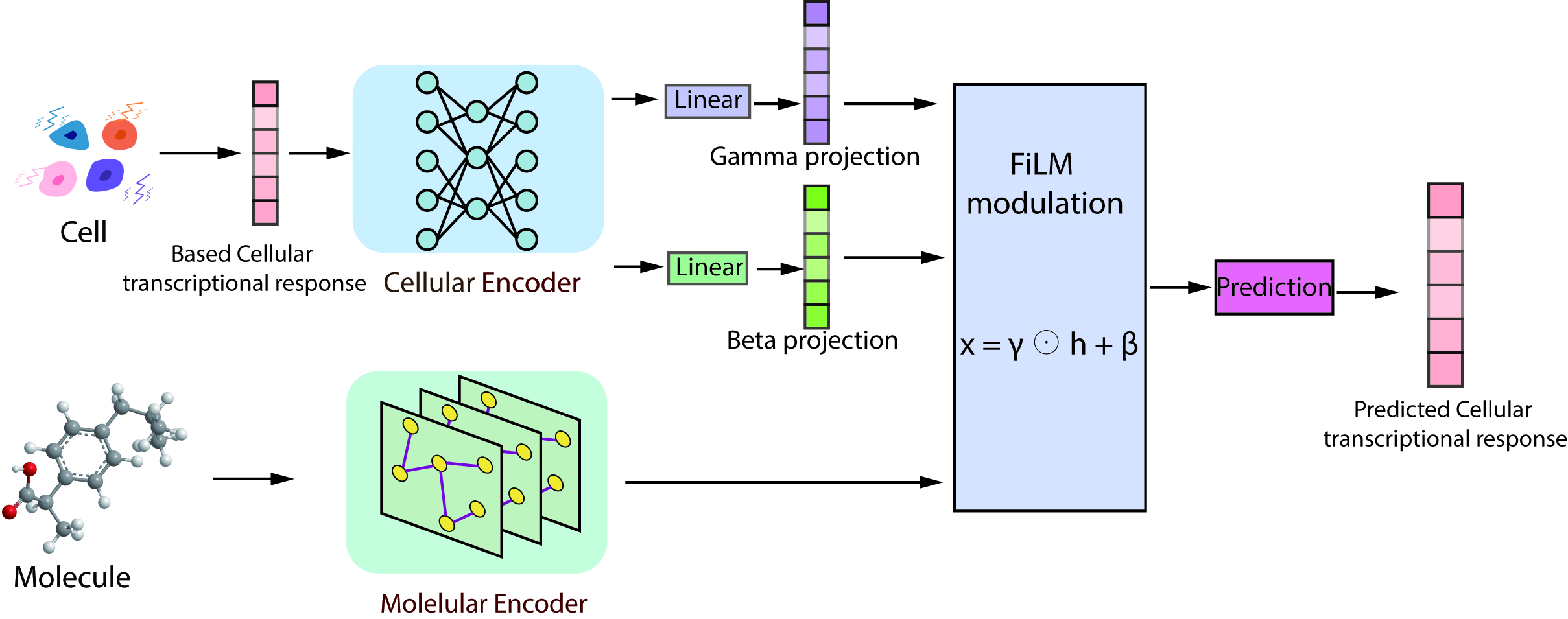

**Figure 6: Architecture of CellChem-Generate.** The model uses a molecule and a basal cellular transcriptional response as inputs. The cellular response is encoded by a cellular encoder and projected into FiLM modulation parameters, γ and β, whereas the molecule is encoded by a molecular encoder. These two branches are integrated through FiLM modulation to generate the predicted cellular transcriptional response.

**Supplementary Tables**

**Table 1: Statistics of small molecule perturbation profiles.**

| **Type** | **Counts** |
| --- | --- |
| Perturbation values | 505052 |
| Perturbation time | 2 |
| Perturbation dose | 15 |
| Cell Type | 20 |

| **Dataset** | **Molecular** | **Protein** | **Interaction** |
| --- | --- | --- | --- |
| Drugbank | 6559 | 4294 | 17512 |
| Clue_drug(seq_based) | 1062 | 884 | 7690 |

**Table 2: Drugbank and CLUE data statistics.**

**Table 3： Evaluation on the Molecule-Protein Interaction Prediction Task with a Randomly Partitioned Test Set.**

| **Methods** | **Accuracy (Std)** | **Precision (Std)** | **Recall (Std)** | **F1 (Std)** | **AUC (Std)** | **AUPR (Std)** |
| --- | --- | --- | --- | --- | --- | --- |
| CellChem | **0.799(0.006)** | **0.768(0.008)** | **0.851(0.023)** | **0.807(0.008)** | **0.864(0.009)** | 0.834(0.016) |
| HyperAttentionDTI | 0.723(0.029) | 0.675(0.032) | 0.850(0.011) | 0.752(0.017) | 0.799(0.023) | 0.783(0.026) |
| TransformerCPI | 0.708(0.011) | 0.703(0.014) | 0.712(0.037) | 0.706(0.016) | 0.765(0.012) | 0.778(0.009) |
| GraphDTA | 0.738(0.011) | 0.771(0.020 | 0.641(0.053) | 0.698(0.024) | 0.773(0.014) | 0.766(0.010) |
| MolTrans | 0.719(0.044) | 0.708(0.072) | 0.776(0.075) | 0.710(0.037) | 0.780(0.020) | 0.762(0.016) |
| GraphBAN | 0.764(0.007) | 0.749(0.011) | 0.759(0.008) | 0.754(0.003) | 0.838(0.005) | 0.834(0.006) |

**Table 4：Ablation Study on Molecular-Protein Interaction Prediction with a Randomly Partitioned Test Set.**

| **Methods** | **Accuracy (Std)** | **Precision (Std)** | **Recall (Std)** | **F1 (Std)** | **AUC (Std)** | **AUPR (Std)** |
| --- | --- | --- | --- | --- | --- | --- |
| CellChem | **0.799(0.006)** | **0.768(0.008)** | **0.851(0.023)** | **0.807(0.008)** | **0.864(0.009)** | **0.834(0.016)** |
| CellChem_wo | 0.785(0.006) | 0.756(0.010) | 0.837(0.011) | 0.794(0.005) | 0.853(0.005) | 0.829(0.012) |

**Table 5：Evaluation on the Molecule-Protein Interaction Prediction Task with a Scaffold Partitioned Test Set.**

| **Methods** | **Accuracy (Std)** | **Precision (Std)** | **Recall (Std)** | **F1 (Std)** | **AUC (Std)** | **AUPR (Std)** |
| --- | --- | --- | --- | --- | --- | --- |
| CellChem | **0.692(0.010)** | **0.713(0.013)** | 0.667(0.046) | **0.689(0.020)** | **0.757(0.009)** | **0.747(0.011)** |
| HyperAttentionDTI | 0.597(0.017) | 0.617(0.019) | 0.556(0.022) | 0.585(0.017) | 0.644(0.023) | 0.640(0.029) |
| TransformerCPI | 0.645(0.010) | 0.670(0.013) | 0.605(0.025) | 0.636(0.014) | 0.702(0.020) | 0.686(0.032) |
| GraphDTA | 0.634(0.029) | 0.645(0.028) | 0.651(0.029) | 0.648(0.028) | 0.682(0.003) | 0.664(0.044) |
| MolTrans | 0.613(0.080) | 0.587(0.061) | 0.933(0.055) | 0.662(0.059) | 0.640(0.131) | 0.645(0.117) |
| GraphBAN | 0.614(0.009) | 0.629(0.012) | 0.600(0.046) | 0.613(0.023) | 0.677(0.008) | 0.667(0.013) |

**Table 6： Ablation Study of Molecular-Protein Interaction Predictions with a Scaffold Partitioned Test Set.**

| **Methods** | **Accuracy (Std)** | **Precision (Std)** | **Recall (Std)** | **F1 (Std)** | **AUC (Std)** |
| --- | --- | --- | --- | --- | --- |
| CellChem | **0.692(0.010)** | **0.713(0.013)** | 0.667(0.046) | 0.689(0.020) | **0.757(0.009)** |
| CellChem_wo | 0.682(0.025) | 0.703(0.011) | 0.655(0.054) | 0.677(0.034) | 0.739(0.020) |

**Table 7: Evaluation of CellChem-Generate in Comparison to Other Methods Using a Random Partition Validation Strategy.**

| **Method** | **Pearson** | **Spearman** | **Positive P@100** | **Positive P@50** | **Positive P@100** | **Negative P@10** | **Negative P@50** | **Negative P@100** |
| --- | --- | --- | --- | --- | --- | --- | --- | --- |
| CellChem | 0.7152 | 0.7107 | 0.8294 | 0.7593 | 0.6955 | 0.8462 | 0.7849 | 0.7290 |
| DeepCE | 0.5156 | 0.5133 | 0.6734 | 0.7216 | 0.6412 | 0.6892 | 0.5861 | 0.6346 |
| CIGER | 0.6965 | 0.6907 | 0.7673 | 0.7302 | 0.6592 | 0.7812 | 0.6884 | 0.7197 |
| MultiDCP | 0.5407 | 0.5394 | 0.4763 | 0.4835 | 0.4389 | 0.4494 | 0.3739 | 0.3947 |
| PRnet | 0.4286 | 0.4299 | 0.86 | 0.52 | 0.708 | 0.656 | 0.596 | 0.666 |
| TransiGen | 0.6797 | 0.7044 | 0.7541 | 0.6192 | 0.5097 | 0.1903 | 0.1865 | 0.1856 |

**Table 8: Evaluation of CellChem-Generate in Comparison to Alternative Methods Using a Cell Type Partition Validation Strategy.**

| **Method** | **Pearson** | **Spearman** | **Positive P@100** | **Positive P@50** | **Positive P@100** | **Negative P@10** | **Negative P@50** | **Negative P@100** |
| --- | --- | --- | --- | --- | --- | --- | --- | --- |
| CellChem | 0.3733 | 0.367 | 0.8597 | 0.5512 | 0.4788 | 0.4342 | 0.5551 | 0.4906 |
| DeepCE | 0.3277 | 0.3243 | 0.4924 | 0.5395 | 0.471 | 0.5159 | 0.4296 | 0.4736 |
| CIGER | 0.3055 | 0.3011 | 0.8597 | 0.4661 | 0.4218 | 0.3837 | 0.5279 | 0.4636 |
| MultiDCP | 0.308 | 0.3036 | 0.3103 | 0.325 | 0.287 | 0.3003 | 0.2481 | 0.2659 |
| PRnet | 0.3259 | 0.3319 | 0.5458 | 0.43 | 0.42 | 0.408 | 0.432 | 0.498 |
| TransiGen | 0.3279 | 0.3250 | 0.9505 | 0.4412 | 0.3375 | 0.2838 | 0.1418 | 0.1411 |

**Table 9: Evaluation of CellChem-Generate in Comparison to Alternative Methods Using a Scaffold Partition Validation Strategy.**

| **Method** | **Pearson** | **Spearman** | **Positive P@100** | **Positive P@50** | **Positive P@100** | **Negative P@10** | **Negative P@50** | **Negative P@100** |
| --- | --- | --- | --- | --- | --- | --- | --- | --- |
| CellChem | 0.3828 | 0.3834 | 0.5262 | 0.4715 | 0.4355 | 0.5612 | 0.5248 | 0.4866 |
| DeepCE | 0.3055 | 0.3079 | 0.4545 | 0.463 | 0.4342 | 0.4951 | 0.3968 | 0.4662 |
| CIGER | 0.3512 | 0.3512 | 0.4771 | 0.455 | 0.402 | 0.5149 | 0.4532 | 0.4575 |
| MultiDCP | 0.2554 | 0.2578 | 0.1938 | 0.2547 | 0.1823 | 0.2457 | 0.16 | 0.2211 |
| PRnet | 0.1755 | 0.1967 | 0.34 | 0.54 | 0.424 | 0.336 | 0.422 | 0.308 |
| TransiGen | 0.3483 | 0.3556 | 0.4121 | 0.3333 | 0.2847 | 0.1599 | 0.1588 | 0.1600 |

**Table 10: Statistics of Functional Diversity and Corresponding Legend for Traditional Chinese Medicine (TCM) Compounds.**

| Effect | Count |
| --- | --- |
| heat-clearing medicinal | 728 |
| blood-activating and stasis-resolving medicinal | 474 |
| heat-clearing and detoxifying medicinal | 473 |
| tonifying and replenishing medicinal | 399 |
| qi-regulating medicinal | 361 |
| exterior-releasing medicinal | 320 |
| interior-warming medicinal | 319 |
| dampness-resolving medicinal | 274 |
| wind-cold-dispersing | 257 |
| hemostatic medicinal | 252 |
| blood-activating analgesic medicinal | 208 |
| cough-suppressing and panting-calming medicinal | 184 |
| blood-activating menstruation-regulating medicinal | 152 |
| wind-dampness dispelling medicinal | 129 |
| qi-tonifying medicinal | 128 |
| water-draining and anti-icteric medicinal | 127 |
| yang-tonifying medicinal | 122 |
| heat-clearing and dampness-drying medicinal | 105 |
| astringent medicinal | 102 |
| meridian-warming hemostatic medicinal | 102 |
